# Indigenous *Bacillus paramycoides* and *Alcaligenes faecalis*: potential solution for the bioremediation of wastewaters

**DOI:** 10.1101/2020.05.20.105940

**Authors:** Aneeba Rashid, Safdar A. Mirza, Ciara Keating, Sikander Ali, Luiza C. Campos

## Abstract

Farmers near towns and cities are using wide range of untreated wastewaters for crop irrigation in Pakistan due to severe freshwater shortage. The present study aimed to treat different types of wastewater including domestic, hospital, textile, pharmaceutical and mixed wastewaters using indigenous bacterial isolates to remove contaminants and render these wastewaters safer for irrigation. 37 bacterial strains were isolated from the 5 wastewater samples collected from different sites in Lahore, Pakistan. Under optimum growth conditions, the isolates D6, D7 and P1 showed maximum decolourisation potential of 96, 96, 93 %, respectively against hospital wastewater. GCMS analysis of the untreated hospital wastewater confirmed the presence of pharmaceutic pollutants i.e. Phenol, Salicylic acid, Caffeine, Naproxen, Octadecene and Diazepam. These organic compounds were biodegraded into derivate Ticlopidine in the case of isolate D6, derivatives Tetradecene and Griseofulvin in the case of isolate D7, and derivatives Lidocaine and Butalbital in the case of isolate P1. 16S rDNA sequencing was used to identify these isolates. Isolates D6 and D7 showed 100 and 99.86 % homology to *Bacillus paramycoides*, a novel strain from *Bacillus cereus* group (Liu et al., 2017). Isolate P1 showed 97.47 % homology to *Alcaligenes faecalis*. These strains therefore could represent a low-cost and low-tech alternative to bioremediate complex wastewaters prior to irrigation to support the achievement of the Sustainable Development Goal 6 - clean water and sanitation in Pakistan.

## 1. Introduction

The Planet Earth contains only less than 1% of freshwater (Gleick, 2014). The increasing population, urbanization, human activities and unjustifiable usage of freshwater are the foremost reasons of causing its further shortage (Khoso et al., 2015). The South Asian region, mainly Pakistan, has the worst condition in this scenario (Roberts, 2017; Wagan and Khoso, 2013). Despite having world’s largest glaciers, researchers have proclaimed that the country is on its way to become the most water-stressed country in the region by year 2040 (WRI, 2015). The country’s agricultural, domestic and industrial sectors have too scored high on the World’s Resource Institute’s water stress index. Its per capita annual water availability is just 1017 m^3^ now (IMF, 2015) which is scarily closer to the scarcity threshold level (1000 m^3^). Being an agricultural country, this scarcity of freshwater resources has driven local farmers in Pakistan to reuse untreated wastewater for irrigation of crops (Mahmood and Malik, 2014). These wastewaters contain many harmful chemicals and heavy metals which accumulate in crops (Afonne and Ifediba, 2020; Topal et al., 2020; Zhang et al., 2020; Zoqi and Doosti, 2020) and up the food chain making them hazardous for consumption.

Pakistan Water Sector Strategy (PWSS, 2002) reported that the total quantity of wastewater produced in Pakistan is 962335 million gal per annum including 674009 million gal from domestic and 288326 million gal from industrial use. The domestic and industrial wastewater is either discharged directly to a sewer system, a natural drain or water body, a nearby field or an internal septic tank in Pakistan (Murtaza and Zia, 2012). Generally, this wastewater is not treated and none of the cities have any biological treatment process except Islamabad and Karachi (EPMS, 2002), and even these cities treat only a small proportion (<8%) of their wastewater before disposal (Bashir, 2012; Steenbergen and Oliemans, 2002). These wastewaters contain considerable amount of dyes, suspended solids, heavy metals, additives, detergents, surfactants, carcinogenic amines and formaldehyde (Azizullah et al., 2011). They also contain organic and inorganic particles and compounds, macro-solids, gases, emulsions, toxins, microplastics (Gatidou et al., 2019), pharmaceuticals like endocrine disrupting compounds, hormones, antibiotics, anesthetics, perfluorinated compounds (Arvaniti and Stasinakis, 2015), siloxanes (Bletsou et al., 2013), drugs of abuse (Gatidou et al., 2016) and various biological pathogens (Andersson et al., 2016). These untreated or insufficiently effluents treated wastewaters pose a serious environmental threat (Salgot et al., 2006). The complex nature of these effluents and lack of centralized wastewater treatment infrastructure make the treatment difficult in Pakistan. One area, that is a considerable challenge is the removal of colour contamination.

The dyes, impurities and chemicals released from the textile industries impart colour to wastewater drains and cause colour contamination, thus diminishing the water quality (Carmen and Daniela, 2012). Various physicochemical methods have been used worldwide to remove colour and impurities from wastewater, i.e. adsorption (Patel and Vashi, 2010), ion exchange (Karcher et al., 2002), membrane filtration (Marcucci et al., 2001), ozonation (Ince and Tezcanli, 2001), photooxidation (Hai et al., 2007) and reverse osmosis (Suksaroj et al., 2005). Pakistan being a developing economy has not adopted any of these methods on a large scale as these methods are prohibitively expensive and require large complex infrastructure (Verma et al., 2012). Only one full scale domestic wastewater treatment plant was set up on the conventional activated sludge process in Islamabad, Pakistan but it is not maintained well by the plant operators (Fatima and Khan, 2012). Decentralised biological treatment methods could offer a potential low-cost and low-tech solution for communities in developing countries such as Pakistan.

Biological processes play a major role in the removal of pollutants. Due to ubiquitous nature of bacteria, they can be used as invaluable tools for the biological treatment of different types of wastewater, i.e. domestic, hospital, pharmaceutical and textile industrial wastewaters. The bioremediation potential of bacterial isolates is an economically viable method and environment friendly thus presents a good alternative to other engineered process (Dwivedi and Tomar, 2018). Biological treatment takes advantage of the catabolic versatility of microorganisms including bacteria to degrade or convert toxic compounds to non-toxic compounds (Díaz, 2008). One strategy – to use native or indigenous isolates from wastewater to degrade, detoxify and decolour specific wastewater has been the source of intensive research. Many authors have isolated microorganisms from industrial textile wastewaters and then demonstrated their ability to decolourise specific classes of dyes in the laboratory (e.g. Zhang et al., 2010; Meerbergen et al., 2018; Alalewi and Jiang, 2012; Buthelezi et al., 2012; Mahmood et al., 2011). Shukor et al (2009) demonstrated isolates from hospital wastewater were capable of degrading acrylamide compounds. Others have used similar strategies to demonstrate the removal of heavy metals (Helmy et al., 2018; Afzal et al., 2017; Das and Kumari, 2016).

Researchers have established the identity of many of these isolates from different wastewaters and their ability to specific chemical compounds, e.g. *Bacillus cereus* isolated from domestic wastewater for degrading acrylamide (Shukor et al., 2009) and hydrocarbons (Kostka et al., 2011), *B. subtilis* isolated from pharmaceutical wastewater for removing antibiotic cephalexin and heavy metals (Adel et al., 2015), *Aeromonas hydrophila* isolated from industrial wastewater for degrading Triarylmethane dyes (Ogugbue and Sawidis, 2011), *Alcaligenes faecalis* spp. isolated from petrochemical industrial wastewater for degrading phenol (Manafi et al., 2011), *Rhodococcus pyridinivorans* isolated from gold mine wastewater for degrading cyanomethane (Sulistinah et al., 2019), *Dracaena sanderiana* isolated from plastic industry wastewater for degrading bisphenol A (Suyamud et al., 2020) or *Sphingomonas trueperi* isolated from wastewater sludge for the degradation of allethrin (Bhatt et al., 2020). However, these biotreatment studies do not represent the complex environment of mixed wastewater. Moreover, isolates are generally tested against specific compounds in simplistic lab conditions and thus, the potential to degrade these compounds in complex raw wastewater is largely unknown. This therefore is not sufficient for the real-world situation in countries like Pakistan where wastewaters from household, hospitals and wide range of industries is combined.

The present research aimed to i) characterise the pollutants and metals in a variety of complex raw wastewaters in Pakistan, ii) isolate novel decolourising isolates from the raw wastewater, iii) determine the decolourisation and degradation potential of these isolates in raw hospital wastewater and finally iv) to identify the isolates with the maximum potential for decolourisation and degradation of organic compounds.

## 2. Materials and methods

### 2.1 Collection of wastewaters

Four wastewater (domestic, hospital, textile and pharmaceutical) samples (50 L each) from the points of discharge of drainage sites in Lahore, Pakistan were collected in sterile bottles according to the standard protocols (APHA, 2005). The geographical coordinates of Lahore city are 31° 34’ 55.3620” north and 74° 19’ 45.7536”’ east at an altitude of 217 m (712 ft). Mixed wastewater (50 L) was also collected from a collective drainage site of the different wastewaters. All five samples were collected in October, 2018. The temperature of the wastewaters and environment were measured on-site with the help of digital thermometer (HUBDIC).

### 2.2 Characterisation of the wastewaters

The wastewaters were analysed immediately after the collection for the characterisation to ensure the bacterial viability and to avoid any self-degradation of organic compounds. Following physicochemical parameters were investigated according to standard protocols (APHA, 2005; Ali et al., 2009) *i.e*. colour, smell, temperature, pH, electrical conductivity (EC), total suspended solids (TSS), total dissolved solids (TDS), chemical oxygen demand (COD), biological oxygen demand (BOD_5_), salinity (ppt) and turbidity (NTU). The concentrations of heavy metals, *i.e*. Arsenic, Cadmium, Chromium, Lead and Nickel, were estimated through Atomic Absorption Spectrophotometer (AA 7000 F with Autosampler and Hydride Vapour Generator, Shimadzu, Japan). The same physicochemical parameters were investigated in treated wastewaters and were compared with untreated wastewaters. Biodegradability index (BI) is the ratio of BOD_5_ : COD. It is a parameter for evaluating the potential biodegradability of a biological treatment in wastewater (Padoley et al., 2012). The values of BI for all decolourised wastewaters were compared with the values of BOD_5_ : COD of the original wastewaters to access the level of biodegradability. The biodegradation of wastewaters with lesser BOD_5_ / COD value is not possible to biodegrade as it contains extremely toxic contaminants. If the BOD_5_ / COD value would be lower than 0.3, then the biodegradation will not proceed, thus it cannot be treated biologically, because the wastewater generated from these activities inhibits the metabolic activity of bacteria due to their toxicity.

### 2.3 Isolation and screening of bacteria

The isolation of bacterial strains from each of the five types of wastewaters was carried out through serial dilution method (Verma et al., 2001). The isolates from each wastewater’s inocula were incubated on sterile nutrient agar medium (0.8% Nutrient broth and 2% Agar) plates in static incubator at 37°C for 24 hours and were then purified by streaking on nutrient agar medium plates. Streaking was done thrice in zig zag manner. The purified cultures were shifted to prepared slants of Luria-Bertani medium (LB) with Agar in test tubes and were preserved in a refrigerator (4 °C) (Mahmood et al., 2011). The bacterial slants were maintained every two weeks on freshly prepared agar slants to circumvent the susceptibility of the isolates (ISO 11133, 2014).

The domestic wastewater was chosen as the preliminary testing sample for screening. The isolated bacterial strains were inoculated (10 %) and incubated at 37 °C for 24 hours in domestic wastewater (100 mL) for initial screening. The percentage decolourisation was measured using UV/VIS (AE-S80) spectrophotometer at 545 nm (Nanthakumar et al., 2013). The bacterial isolates showing more than 50% decolourisation were then tested and inoculated using the same methodology against each type of wastewater separately (D5, D6, D7, D8, H6, T4, T5, T6, P1, M5 and M8). The bacterial isolates showing maximum decolourisation (more than 90%) against all wastewaters tested were further selected for testing optimal conditions (See supplementary data) for colour contamination removal in complex wastewaters.

### 2.4 Testing decolourisation potential of isolated bacteria

The parameters incubation time, temperature and inoculum concentration were selected for the estimation of optimal growth conditions of three bacterial isolates for testing their decolourisation potential. For incubation time, the conical flasks (250 mL) containing 100 mL of domestic wastewater each were inoculated with the screened isolates (10 % inoculum) in shaking incubator at 120 rpm (PMI Labortechnik GMBH, WIS-20R) (Taran et al., 2007). The flasks were incubated for 24, 48, 72 and 96 hours at 37 °C. For testing the optimal temperature, the inoculum (10%) of screened isolates was added to domestic wastewater (100 mL) in conical flasks (250 mL) for 24 h. Flasks were incubated at 30, 37, 44, 51 and 58°C in a shaking incubator (PMI Labortechnik GMBH, WIS-20R). For the inoculum concentration, a loop full of bacterial colony from a plate was added in distilled water (100 mL). The optical density (OD) was adjusted to 1 at 545 nm wavelength using UV/VIS spectrophotometer (A&E Labmed, AE-S80) in order to maintain equal number of bacterial cells to each inoculum. The inoculum concentrations tested were 5, 10, 15, 20, 25 and 30 % (Getha et al., 1998). The bacterial cell count per mL of each screened isolate was also done through haemocytometer slide bridge (Neubaur improved HBG, Marinefield, Germany). On optimum inoculum concentration (10 %), optimum incubation time (48 h) and optimum temperature (37 and 51 °C), the decolourisation tests was conducted.

The three bacterial isolates were inoculated (10 %) separately in five types of wastewaters (100 mL each) present in conical flasks (250 mL) for 48 hours at 37 and 51 °C (Jadhav et al., 2010). The percentage decolourisation was calculated using Equation (1) (Cheriaa et al., 2012) at 545 nm using UV/VIS (AE-S80) spectrophotometer:

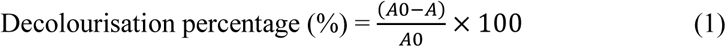

Where A_0_= Initial absorbance, A= Absorbance of medium after decolourisation at the λ_max_ (nm). The decolourisation experiments were performed in triplicates.

#### 2.4.1 Organic compounds degradation

Bioremediation potential of the hospital wastewater sample that showed maximum decolourisation percentual and biodegradability index (Section 2.2 and 2.4) was further analysed for organic compounds degradation. The hospital wastewater sample was analyzed by gas chromatography mass spectrometry (GCMS) technique using an Agilent Gas Chromatograph (GC, AgiTech-7260) and Mass Spectrometer (MS, Maspec-6595). In total, four samples (10 mL each) were prepared for the analysis, *i.e*. one uninoculated hospital wastewater sample (control) and three inoculated (i.e. decolourised) hospital wastewater samples. The inoculated three samples were centrifuged (8000g for 15 min) to remove the biomass and the supernatants were shifted in polypropylene falcon tubes (15 mL). All the samples were acidified to pH 1–2 with concentrated HCl and then thoroughly extracted with three volumes of ethyl acetate. The organic layer was collected, dewatered over anhydrous Na_2_SO_4_ and filtered through Whatman filter paper (no. 54). All the GC separations were accomplished using a 20 m×0.3 mm (as internal diameter) fused-silica capillary column with a 0.45 μm coated 6% phenylmethyl silicone film in the instrument.

The aliquot of the sample (5 μL) was injected in split-less mode (0.5 min) at 290°C. The oven temperature was set as follows: initial temperature (45°C), raised to 58-92°C/min and then 12-210°C/min, 10-285°C/min and 6-320°C/min with a hold time of 5 min. The pressure control was adjusted for a constant electronic flow of helium as the carrier gas (mL/min). Mass Spectrometer was adjusted as follows: 120°C analyzer, 210°C source, 280°C interface and electron ionization at 80 eV. The data was collected from 50-450 atomic mass unit (amu). The retention time (±0.1 min), quantification ions, confirmation ions (156.18 and 184.25 m/z) and internal standards (Acenaphthene and Phenanthrene) of each sample were set at optimal levels (Spiking level = 0.05 µg/g; recovery = 98.9 and 93.47 %; coefficient of variation (CV) = 4.22 and 7.39 %) and run in accordance with the system sequence. The base-peak ion was employed for quantitation and two qualifier ions were used for confirmation. The compound concentrations were compared with internal standard quantitation (LoQ = 0.05 mg/kg) and calibration curves. In order to identify the low molecular weight compounds derived from bacterial treatment, the mass spectra were compared with National Institute of Standard and Technology (NIST) database library software available in the instrument and by comparing the retention time with those of authentic compounds available. Quantification of these compounds was conducted by relating the ratio of the peak area of the compound of interest over the peak area of the internal standard (Acenaphthene and Phenanthrene) to the calibration curve of standard solution.

### 2.5 Metal tolerance limits

100 mg/L solutions of following ten metal salts were prepared in deionized water, *i.e*. PbNO_3_, CoCl_2_, CaCl_2_, ZnSO_4_, MnSO_4_, MgSO_4_, FeSO_4_, K_2_Cr_2_O_7_, Na_2_MoO_4_ and CuSO_4_ (Sigma Aldrich, Uk). The three bacterial isolates were streaked on the prepared metal salt-nutrient agar plates and kept in static incubator (at 37°C) for 24 h. At 100 mg/L concentration, the metal salt plates with more than 65% bacterial growth were selected. The solutions of these metal salts (CaCl_2_, MgSO_4_, K_2_Cr_2_O_7_, Na_2_MoO_4_ and PbNO_3_) were then prepared in 50, 100, 150, 200, 250 and 300 mg/L concentrations. The bacterial growth of three isolates on these concentrations was assessed.

### 2.6 Identification of the bacterial isolates

The three bacterial isolates showing > 90% decolourisation potential above were selected for identification through 16S rDNA sequencing (Mignard and Flandrois, 2006). Neat DNA (0.5 mL) was sent to the Macrogen sequencing company in South Korea for sequencing analysis. Polymerase chain reaction (PCR) was carried on the three isolates using the following forward and reverse primer set (See supplementary data): 27F (AGA GTT TGA TCM TGG CTC AG) and 1492R (TAC GGY TAC CTT GTT ACG ACT T) (Muyzer et al., 1993). 20 ng of genomic DNA template was taken in a 30 µL reaction mixture using *EF-Taq* (SolGent, Korea) as follows: *Taq* polymerase activation for 2 min at 95°C, 35 cycles for 1 min at 95°C, 1 min each at 55°C and 72°C were performed finishing with 10 min step at 72°C. Amplification products were purified with a multiscreen filter plate (Millipore Corporation, Bedford, Ma, USA). The sequencing reaction was performed using a PRISM BigDye Terminator v3.1 Cycle sequencing Kit. DNA samples containing the extension products were added to Hi-Di formamide (Applied Biosystems, Foster City, CA). The mixture was incubated for 5 min at 95°C, followed by 5 min on ice and then analyzed by ABI Prism 3730XL DNA analyzer (Applied Biosystems, Foster City, CA).

The forward and reverse sequence chromatograms (abi files) were initially viewed in FinchTV version 1.5.0 and then interrogated using MacVector version 17.5.4. Raw sequences were examined in MacVector and ambiguous bases were edited by comparing the individual electrograms per strain. Low quality ends were trimmed. The forward and reverse reads were imported into BioEdiT version 7.2. A consensus sequence per strain was subsequently assembled using the contig assembler program (CAP; Huang, 1992) using the forward read and reverse complement of the reverse read. The full sequence information and raw chromatogram details are presented in the Supplementary Information. BLAST analysis was carried out on the assembled sequences. The sequences of the three isolates were deposited in GenBank with accession numbers. Phylogenetic analysis of the strains was carried out using the top 20 BLAST hits for each isolate. This was achieved by aligning the sequences using Muscle version 3.8.425 (Edgar, 2004) and a phylogenetic tree assembled in Geneious Prime using Tamura-Nei genetic distance method and Neighbor-Joining tree building method. This tree was then imported in newick file format and edited in Evolview (Zhang et al., 2012).

## 3. Results and discussion

### 3.1 Characterisation of the wastewaters

The apparent colours of domestic, hospital, textile, pharmaceutical and mixed wastewaters were light grey, light yellow, greenish grey, light brown and blackish, respectively. The true colour values for the wastewaters were 101, 188, 221, 103 and 311 PCU, respectively. The smell of the domestic, textile and mixed wastewaters was pungent, while hospital and pharmaceutical wastewaters had fishy smell. The values of most of the physicochemical parameters were beyond the level of National Environment Quality Standards (NEQS, 2000). Like the pH values of textile and pharmaceutical wastewater were 8.7 and 10.4, respectively, before treatment keeping in mind that the NEQS range for pH is 6.6-8.5. The pH values of domestic, hospital and mixed wastewaters were within the NEQs range before treatment. This indicated that these wastewaters had their pH corrected before being discharged. Similarly, the values of total suspended solids of domestic, hospital, textile, pharmaceutical and mixed wastewaters were 1920, 2300, 2150, 2120 and 2670 mg/L. These values were too beyond the range of NEQS standard (i.e. < 500 mg/L). The values for the turbidity should be less than 5 NTU as per NEQS range. While the turbidity values for domestic, hospital, textile, pharmaceutical and mixed wastewaters were 38, 51, 76, 61 and 123 NTU (Nephelometric Turbidity Units), respectively.

The wastewaters became colourless and odourless after the biotreatment (Figure 1 a,b). The true colour values for domestic, hospital, textile, pharmaceutical and mixed wastewaters were reduced to 28, 55, 61, 38 and 64 PCU, respectively. Results showed that the values of analyses of various physicochemical parameters were within the levels of National Environment Quality Standards (NEQS, 2000) after treatment (Table 1). Like, the pH of textile and pharmaceutical wastewaters after the treatment were reduced to 7.5 and 8.2, respectively (pH range: 6.6-8.5). The reduction in pH after the decolourisation of textile wastewater has been reported previously by Ogugbue and Sawidis (2011b). Similarly, the values of total suspended solids (TSS) were reduced to 363, 483, 425, 398 and 491 mg/L after the biotreatment (TSS range: < 500 mg/L). The turbidity values were reduced after the biotreatment to 4, 5, 5, 3 and 4 which were within the range of NEQS turbidity value (i.e. ≤ 5 NTU).

**Table 1:**
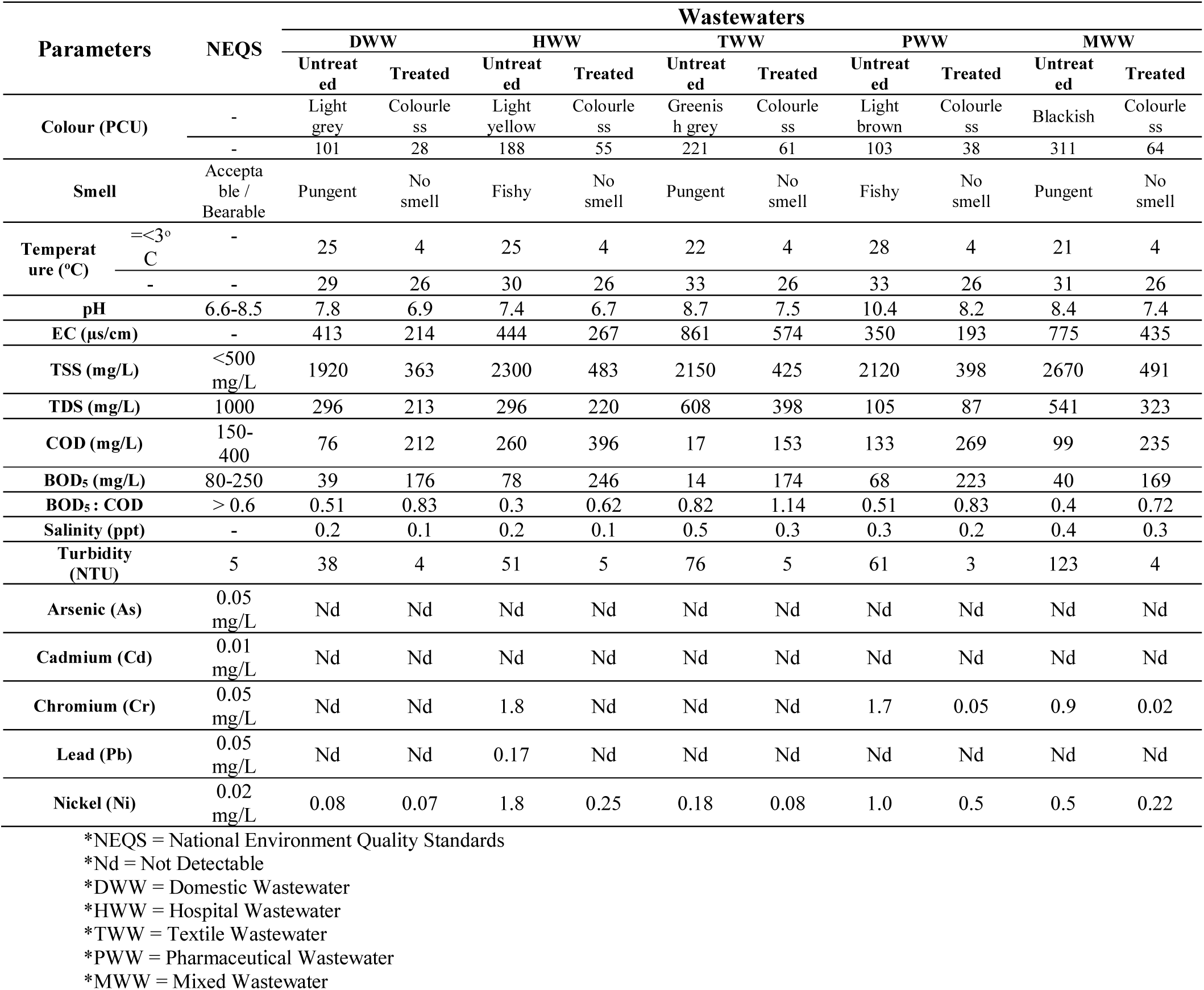
Physiochemical characterisation of untreated and treated wastewaters in comparison to NEQS

**Figure 1:**
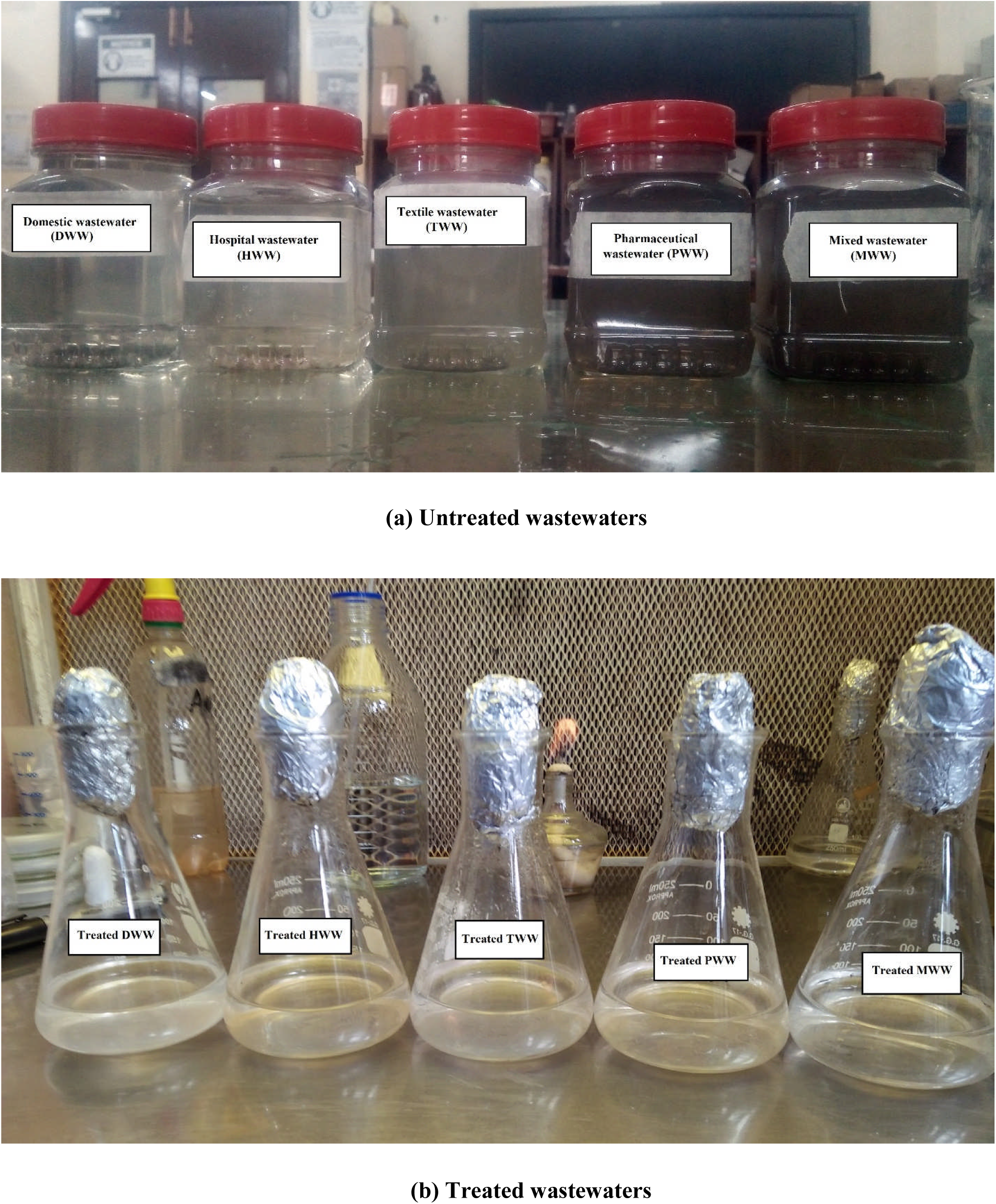
Comparison of wastewaters before and after decolourisation.

The values of BOD_5_ for domestic, hospital, textile, pharmaceutical and mixed wastewaters were 39, 78, 14, 68 and 40 mg/L, respectively. This was out of the range of NEQs which is 80 – 250 mg/L. The values of COD for these wastewaters were 76, 260, 17, 133 and 99 mg/L, respectively. The value of COD in all wastewaters were below the range (150 – 400 mg/L) except hospital wastewater. The BOD_5_ / COD ratio for these wastewaters were 0.51, 0.3, 0.82, 0.51 and 0.4, respectively. One important thing to notice is that if the value of BOD_5_ / COD is in between 0.3 and 0.6, then wastewater is required to treat it biologically, because the process would be relatively slow, as the acclimatization of the microorganisms that help in the degradation process takes time (Abdalla and Hammam, 2014). All of our wastewater samples lie in the same range between 0.3 and 0.6. However, the lowest value of this ratio recorded was of hospital wastewater that showed it was the most contaminated wastewater than all other types.

The values of BOD_5_ after the biotreatment of domestic, hospital, textile, pharmaceutical and mixed wastewaters were 176, 246, 174, 223 and 169 mg/L, respectively that were within the range of NEQs (80 – 250 mg/L). The values of COD for these wastewaters after biotreatment were 212, 396, 153, 269 and 235 mg/L, respectively that were too within the range (150 – 400 mg/L). The BOD_5_ / COD ratio for these wastewaters were 0.83, 0.62, 1.14, 0.83 and 0.72, respectively. As per previously reported work, the value of BOD_5_ / COD ration > 0.6 confirms the biotreatment of wastewater (Abdalla and Hammam, 2014). All the values of biodegradability index in our wastewater samples were more than 0.6. Even the value of most contaminated hospital wastewater was also 0.62 that showed significant biodegradability index.

Heavy metal chromium was detected in the hospital (1.8 mg/L), pharmaceutical (1.7 mg/L) and mixed (0.9 mg/L) wastewaters which was exceeding the NEQs limit (< 0.05 mg/L). Lead was only present in the hospital wastewater (0.17 mg/L). Nickel was present in domestic (0.08 mg/L), hospital (1.76 mg/L), textile (0.19 mg/L), pharmaceutical (1 mg/L) and mixed (0.5 mg/L) wastewaters (Table 1). The hospital wastewater seemed to have more heavy metals than all other types of wastewaters under study. After treatment, the chromium became absent in hospital wastewater. Its amount was reduced to the NEQ limit (< 0.05 mg/L) in pharmaceutical (0.05 mg/L) and mixed (0.019 mg/L) wastewaters after biotreatment (Table 1). Lead which was only present in the hospital wastewater was not detected after biotreatment. The values of Nickel were reduced to 0.07, 0.25, 0.08, 0.5 and 0.22 after the biotreatment of domestic, hospital, textile, pharmaceutical and mixed wastewaters, respectively. Our results agree well with previous work. For example, Abo-Amer et al. (2015) and Naik et al., (2012) have reported the removal of heavy metals from sewage and electroplating wastewaters, respectively. Also, Ali et al. (2009) have reported reduction in colour, temperature, pH, EC, BOD_5_, COD, TSS, TDS and heavy metals ions present in textile wastewaters after the bioremediation by isolated bacteria.

#### 3.2 Isolation and screening of bacteria

In total, 37 bacterial strains were isolated from domestic, hospital, textile, pharmaceutical and mixed wastewaters. Eight bacteria were isolated from the domestic wastewater (D1-D8), nine bacteria were isolated from the hospital wastewater (H1-H9), six from the textile wastewater (T1-T6), six from the pharmaceutical wastewater (P1-P6) and eight were isolated from the mixed wastewater (M1-M8). The isolations of bacteria have been reported from domestic (Jin et al., 2015), hospital (Yamina et al., 2014), textile (Alalewi and Jiang, 2012) and pharmaceutical (Madukasi et al., 2010) wastewaters. Meerbergen et al. (2018) isolated the bacterial isolates from textile wastewater to decolourise azo dyes. Similarly, four bacterial strains were isolated from marine and tannery saline wastewater samples that were proven to be salt-tolerant and carried out successful bioremediation (Sivaprakasam et al. 2008). Shomar et al. (2020) researched on the significance of using the isolated (viable) bacteria for wastewater treatments.

Eleven bacteria, isolated from domestic, hospital, textile, pharmaceutical and mixed wastewaters, had the potential to decolourise the preliminary tested domestic wastewater in comparison with other bacterial isolates under study. The percentage decolourisations of these bacterial strains isolated from domestic (D5, D6, D7 and D8), hospital (H6), textile (T4, T5, and T6), pharmaceutical (P1) and mixed wastewaters (M5 and M8) were > 50% (Figure 2). After final screening, three bacterial strains showed more than 70% decolourisation potential against all wastewaters *i.e*. D6, D7 and P1 (Figure 3). The isolate D6 exhibited 71, 93, 70, 83 and 73 % decolourisation of domestic, hospital, textile, pharmaceutical and mixed wastewaters, respectively. The isolate D7 showed 74, 91, 70, 83 and 73 % decolourisation of domestic, hospital, textile, pharmaceutical and mixed wastewaters, respectively. The isolate P1 showed 82, 92, 71, 77 and 75 % decolourisation of domestic, hospital, textile, pharmaceutical and mixed wastewaters, respectively. Chen et al. (2003) reported varied decolourisation capabilities (14 – 90 %) of six bacterial strains isolated from textile wastewater for azo, anthraquinone and indigoid dye groups. Meerbergen et al. (2018) reported > 80 % decolourisation potential of five bacterial strains isolated from domestic wastewater treatment plant to decolourise azo dyes. However, most of the work has been done on synthetic components of textile wastewaters (e.g. azo dyes) while our work has provided a complex combination and is more representative of the real-world scenario in Pakistan.

**Figure 2.**
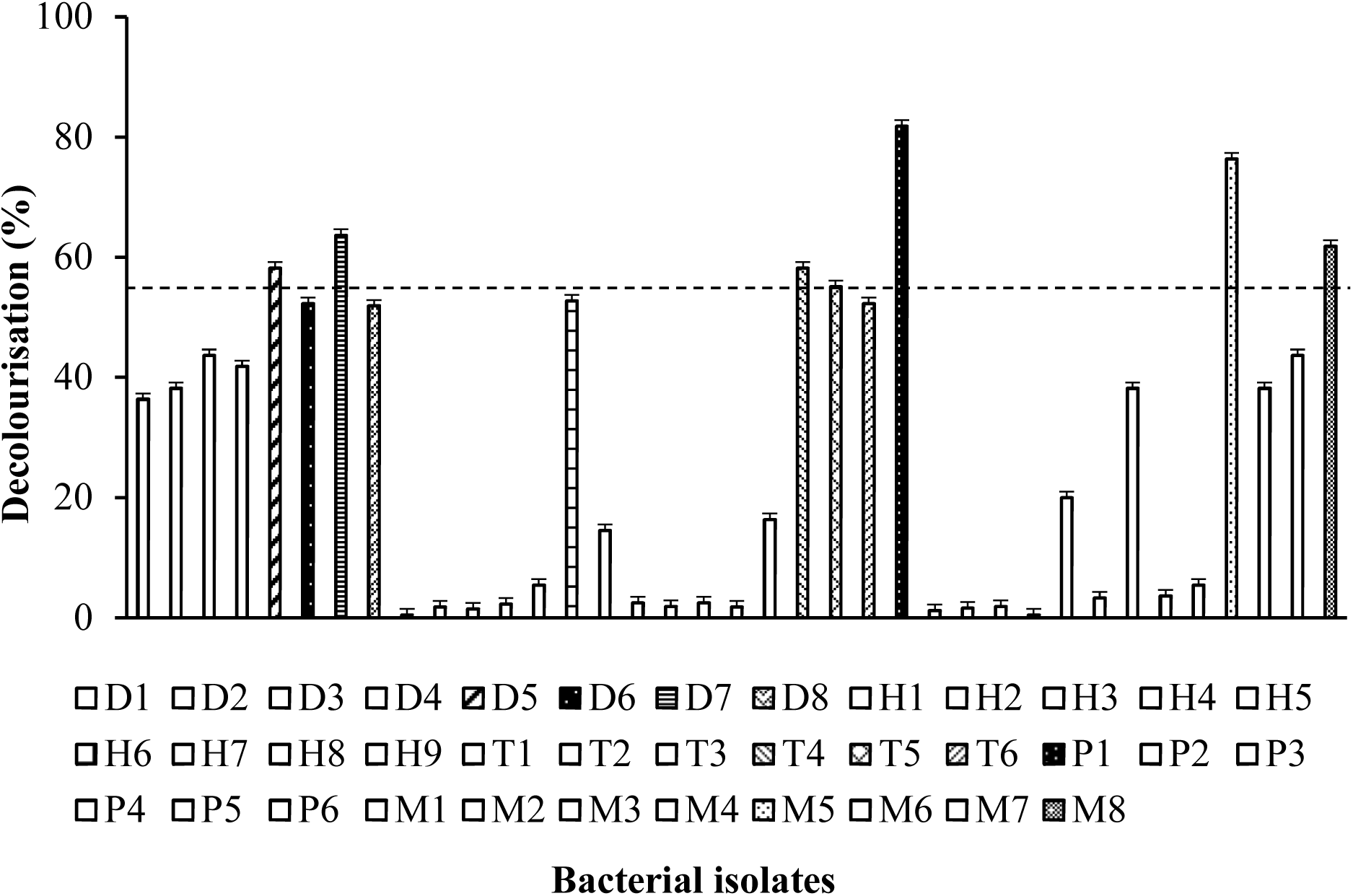
Initial screening from 37 bacterial isolates (>50%); D1-D8 in domestic wastewater, H1-H9 in hospital wastewater, T1-T6 in textile wastewater, P1-P6 in pharmaceutical wastewater and M1-M8 in mixed wastewater

**Figure 3.**
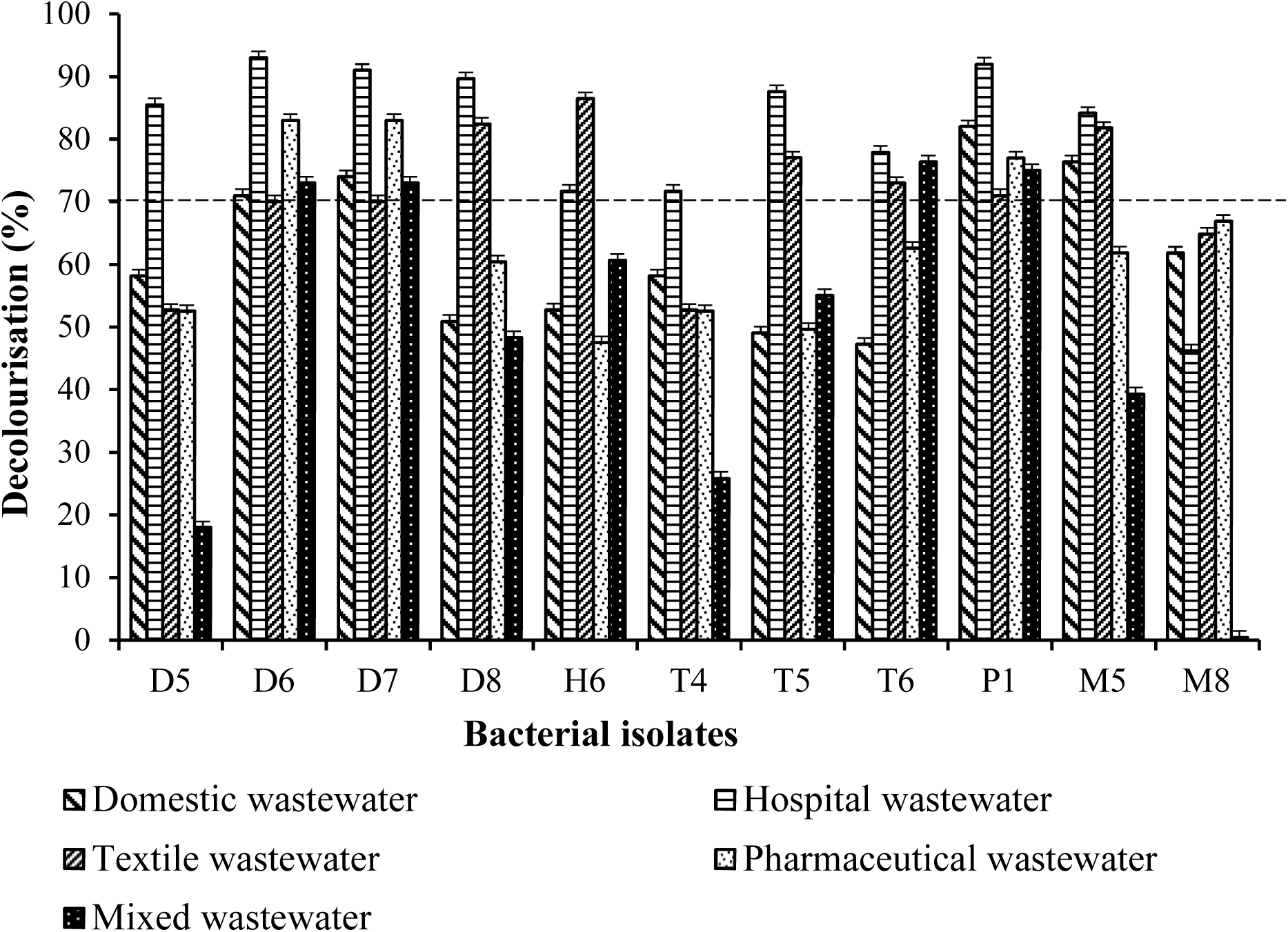
Final screening from 11 bacterial isolates (>70%) against domestic, hospital, textile, pharmaceutical and mixed wastewaters

### 3.3 Testing decolourisation potential of isolated bacteria

The strain D6 exhibited 87, 96, 80, 93 and 83 % decolourisation of domestic, hospital, textile, pharmaceutical and mixed wastewaters, respectively. The strain D7 showed 84, 96, 88, 89 and 83 % decolourisation of domestic, hospital, textile, pharmaceutical and mixed wastewaters, respectively. The strain P1 showed 89, 93, 81, 87 and 85 % decolourisation of domestic, hospital, textile, pharmaceutical and mixed wastewaters, respectively (Figure 4). The high decolourisation potential of 95-98 % have been reported previously in textile wastewater (Deng et al. 2008). Similarly, Saha et al. (2017), Modi et al. (2010), Kanagaraj et al. (2012) and Liao et al. (2013) have also worked on the decolourisation potential of bacterial isolates for textile wastewater. However, to our knowledge, this study has proven significant regarding the decolourisation and bioremediation potential of these strains for pharmaceutical industrial, hospital, domestic and mixed wastewaters that are frequently discharged in Pakistan.

**Figure 4.**
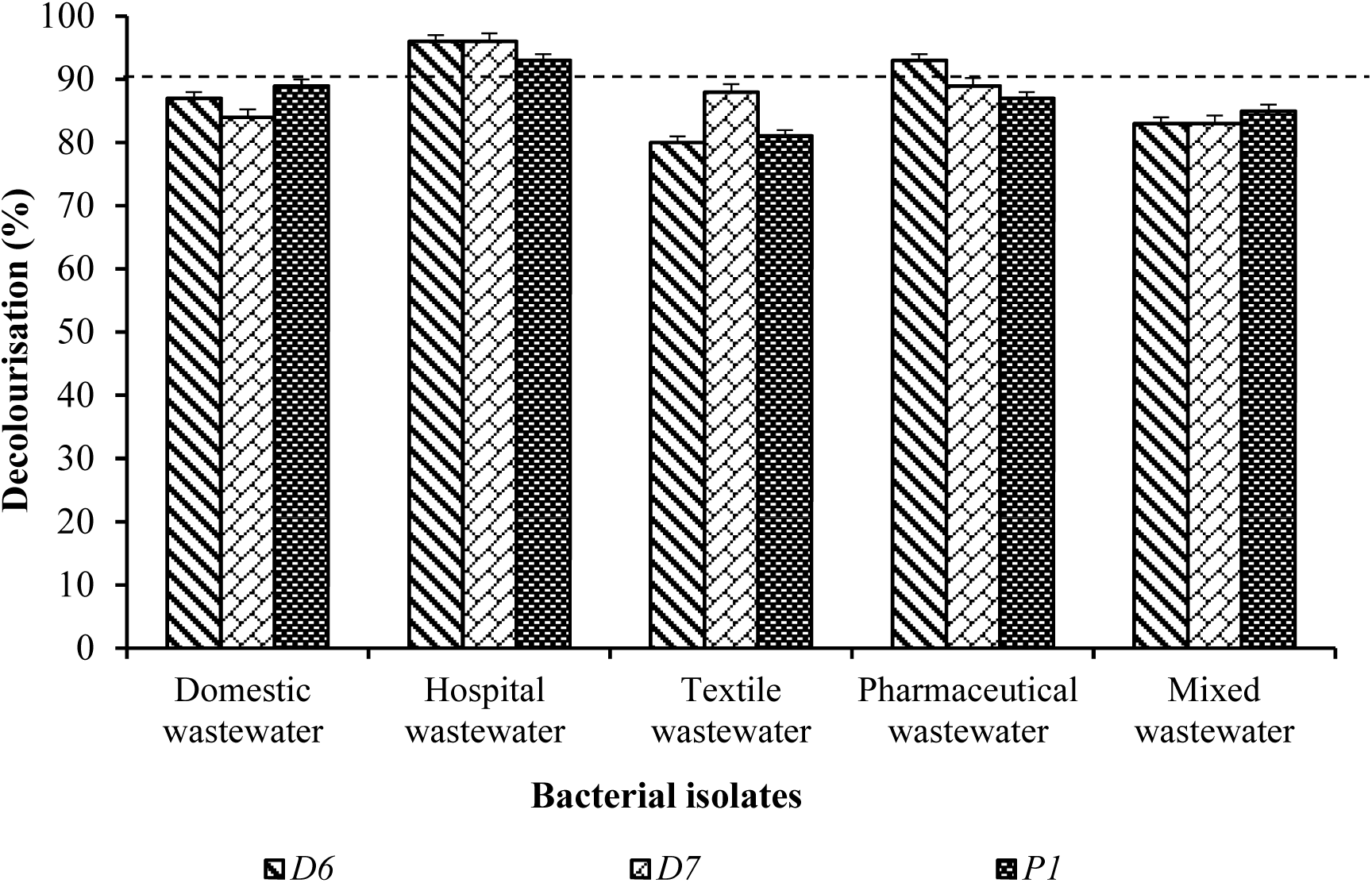
Decolourisation of domestic, hospital, textile, pharmaceutical and mixed wastewaters against D6, D7 and P1

#### 3.3.1 Organic compounds degradation

Considering the maximum decolourisation potential, fluctuating physicochemical values and significant biodegradability index value against bacterial isolates D6, D7 and P1, the untreated and decolourised samples of hospital wastewater were analyzed for degradation of organic compounds. GCMS analysis of untreated hospital wastewater confirmed the presence of six pharmaceutic pollutants in the effluent. These pollutants belonged to following different major groups: aromatic, metabolite, stimulant, NSAID, organic and sedative (Table 2). The pollutants belonging to these groups (with concentrations) were Phenol (0.876 ppm), Salicylic acid (0.048 ppm), Caffeine (0.007 ppm), Naproxen (0.023 ppm), Octadecene (0.185 ppm) and Diazepam (0.014 ppm). The retention time (min) for these pollutants were 26.72, 6.51, 7.96, 9.16, 28.65 and 38.06 minutes, respectively. The confirmation (m/z) ion for these pollutants were 58.15, 147.64, 266.82, 412.07, 581.46 and 685.39 m/z, respectively (See supplementary data). Nair et al., (2008) have described the hazardous nature of phenolic pollutants even at relatively low concentration. Accumulation of phenol creates toxicity both for flora and fauna. Rodil et al., (2012) have reported that salicylic acid is one of the emerging most concentrated pollutant (exceeding the 1 μg/L) which is very hard to remove from the wastewaters even after biotreatment. Motuzas et al. (2017) have reported caffeine as an environmentally emerging micro-pollutant. The presence of non-steroidal anti-inflammatory drug (NSAID) like naproxen in the environment is an emerging problem due to their potential influence on human health and biocenosis or microbial communities (Wojcieszynska et al., 2014). Octadecene was found as an organic priority pollutant in Potato crop (concentration = 0.06 mg/kg; retention time = 21.12 minutes) that was irrigated with ground water having pesticides and herbicides residues (Gushit et al., 2013). Rosal et al. (2010) reported diazepam as an emerging pollutant in urban wastewater with an average concentration of 3 ng/L (that equals LOQ).

**Table 2:**
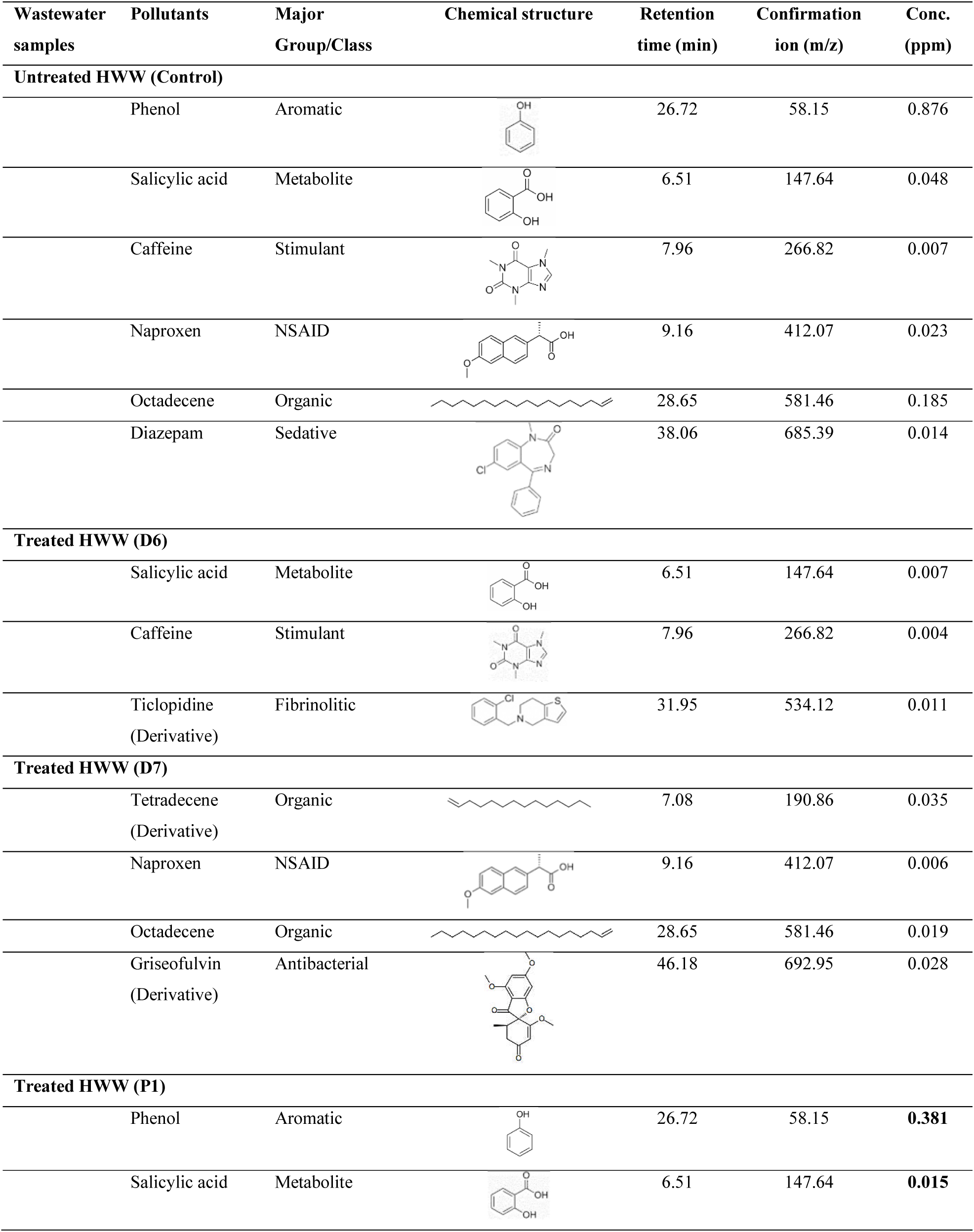
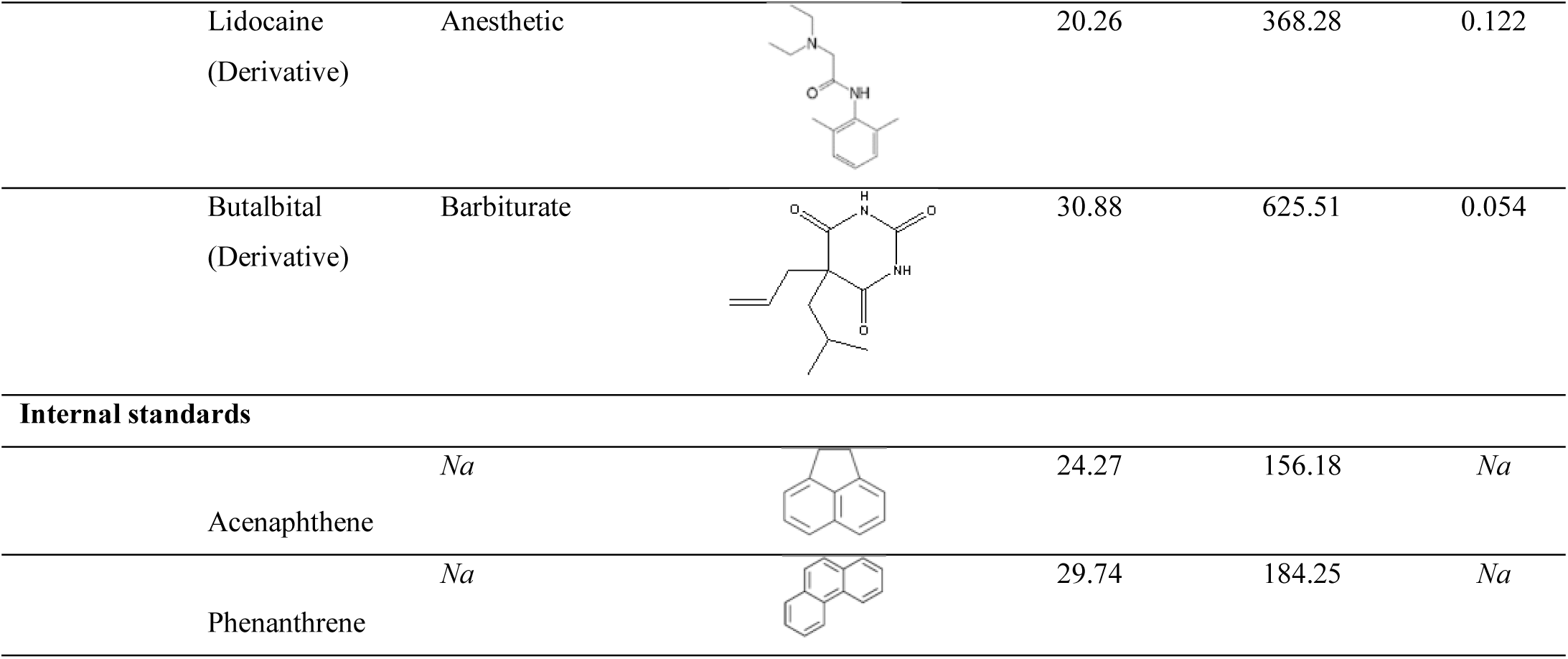
Analysis of untreated and treated hospital wastewater through GCMS.

In the hospital wastewater sample treated with bacterial isolate D6, all other four pollutants were completely biodegraded except salicylic acid and caffeine which were now present at very low concentrations (0.007 and 0.004 ppm) that showed their partial degradation leading to reduction in its concentration from its concentrations in untreated sample 0.048 and 0.007 ppm, respectively. However, a new intermediate compound Triclopidine belonging to Fibrinolitic group was found with 0.011 ppm concentration, 31.95 minutes retention time and 534.12 m/z confirmation ion. Previous researches have supported our ecofriendly biodegradation in this treated sample as Ticlopidine helps in prevention of stroke even better than Aspirin (Grotta et al., 1992). It is also helpful in coronary stenting and as antiplatelet agent during coronary interventions to cure the patients with acute myocardial infarction (AMI) (Cherian et al., 1998).

In the hospital wastewater sample treated with bacterial isolate D7, all other four pollutants were completely biodegraded except naproxen and octadecene which were now present at very low concentrations (0.006 and 0.019 ppm) that showed their partial degradation leading to reduction in its concentration from its concentrations in untreated sample 0.023 and 0.185 ppm, respectively. However, two new intermediate compounds Tetradecene and Griseofulvin belonging to organic and antibacterial groups were found present with 0.035 and 0.028 ppm concentrations, 7.08 and 46.18 minutes retention times and 190.86 and 692.95 m/z confirmation ions. The formation of these essentially important compounds has been supported by previous researches. For example, Roth et al. (1959) reported the Griseofulvin as an antifungal and antibiotic. It is very interesting that a bacterial strain has helped in the formation of an antibiotic through degradation of organic pollutants. Similarly, Tetradecene is a very important compound used in making polyalphaolefins (PAO) at a very low viscosity and excellent cold temperatures (Goze et al., 2007).

In the hospital wastewater sample treated with bacterial isolate P1, all other four pollutants were completely biodegraded except phenol and salicylic acid (0.381 and 0.015 ppm). However, two new intermediate compounds Lidocaine and Butalbital belonging to anesthetic and barbiturate groups were found present with 0.122 and 0.054 ppm concentrations, 20.26 and 30.88 minutes retention times and 368.27 and 625.51 m/z confirmation ions. Previously reported work has supported this biodegradation as an ecofriendly one. For example, Lidocaine is said to possess analgesic (Hollmann et al., 2000; Hollmann et al., 2005), antihyperalgesic (Nagy et al., 1996) and anti-inflammatory (Sugimoto et al., 2003) properties. It is also known to accelerate the return of bowel function after surgery (Marret et al., 2008). It is helpful for post-operative pain and acute rehabilitation after laparoscopic nephrectomy (Tauzin-Fin et al., 2014). Additionally, Butalbital is an analgesic usually prescribed for the treatment of migraine and tension-type headaches (Silberstein and McCrory, 2001). The maternal periconceptional use of butalbital also supports in healing congenital heart defects (Browne et al., 2013). However, its overuse causes headache and discontinuation syndromes (Devine et al., 2005).

### 3.4 Metal tolerance limits

At 100 mg/L concentration of metal salts of PbNO_3_, MgSO_4_, MnSO_4_, ZnSO_4_, K_2_Cr_2_O_7_, CaCl_2_, Na_2_MoO_4_, CuSO_4_, CoCl_2_ and FeSO_4_, the isolate D6 exhibited growth of 25, 70, 50, 5, 35, 80, 45, 20, 5 and 50 %, respectively. It showed maximum growth of 80 % against CaCl_2_. The isolate D7 indicated growth of 65, 35, 25, 12, 20, 35, 45, 3, 0 and 45%, respectively. It showed maximum growth of 65 % against PbNO_3_. The isolate P1 showed growth of 95, 65, 40, 15, 60, 75, 95, 20, 0 and 35 %, respectively. It showed maximum growth of 95 % against both PbNO_3_ and Na_2_MoO_4_ (Figure 5).

**Figure 5.**
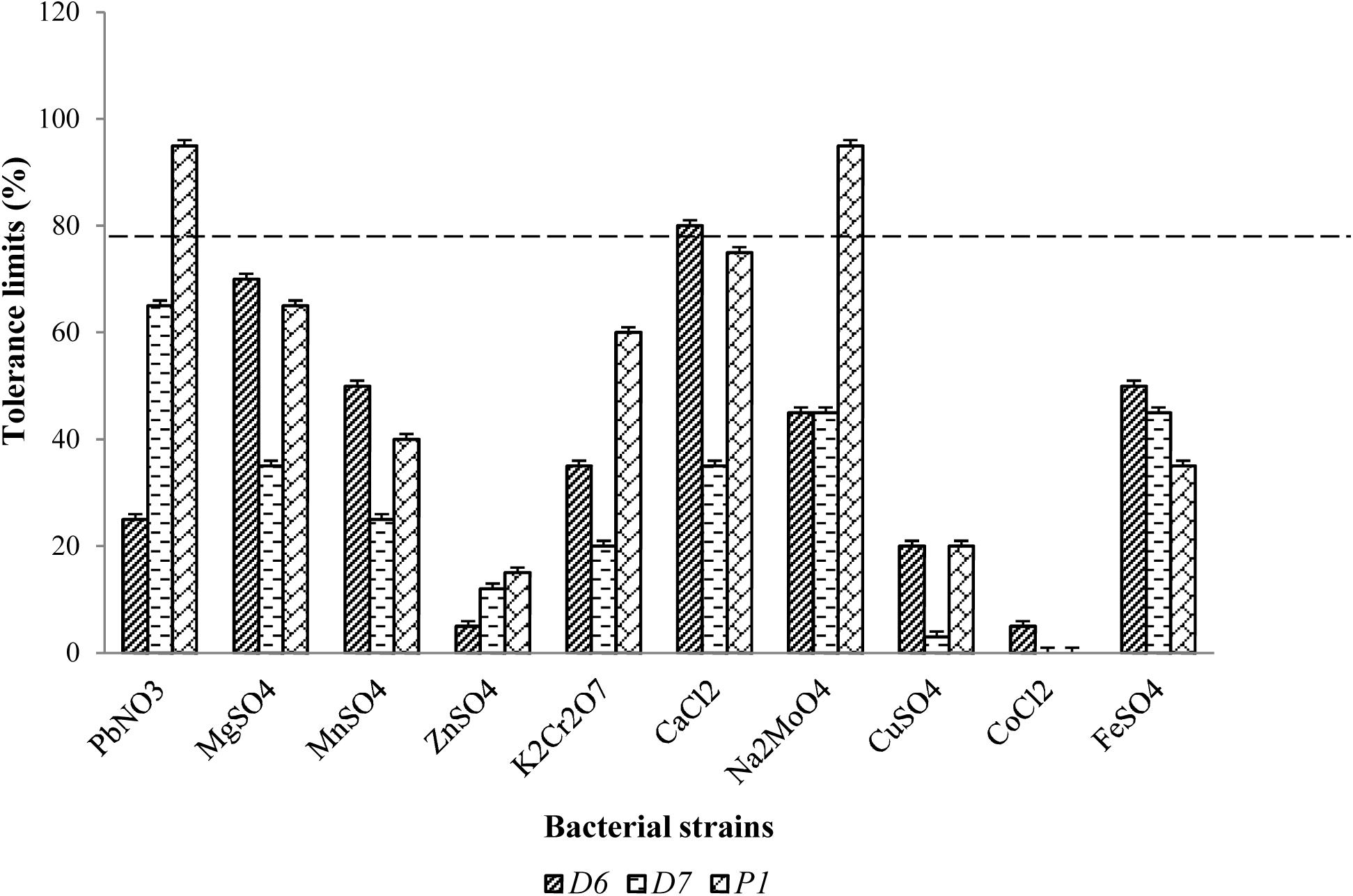
Tolerance limits of bacterial isolates against metal salts (>65%)

For isolate D6, CaCl_2_ and MgSO_4_ metal salts were selected that showed overall maximum growth of 78 and 70 % at 300mg/L concentrations, respectively. For isolate D7, PbNO_3_ metal salt was selected that showed maximum growth of 82 % at 300mg/L concentration. For isolate P1, PbNO_3_, Na_2_MoO_4_, CaCl_2_, MgSO_4_ and K_2_Cr_2_O_7_ metal salts were selected that showed maximum growth of 65, 90, 73, 73 and 75 % at 300mg/L concentration of these metal salts, respectively (Figure 6 a,b,c). On one hand, this has confirmed that all three strains have the potential to tolerate these metals efficiently along with remediating the organic compounds from wastewaters even in co-existence with heavy metals. On the other hand, it also supported our results (Section 3.1) that these isolates have potential to adsorb the heavy metals to remove them from wastewaters. The high metals concentration is really a big challenge for wastewater treatments as it leads to the inhibition of the microbial populations etc. These strains were resistant to high metal concentrations and thus tolerated the harsh environments of these complex wastewaters.

**Figure 6.**
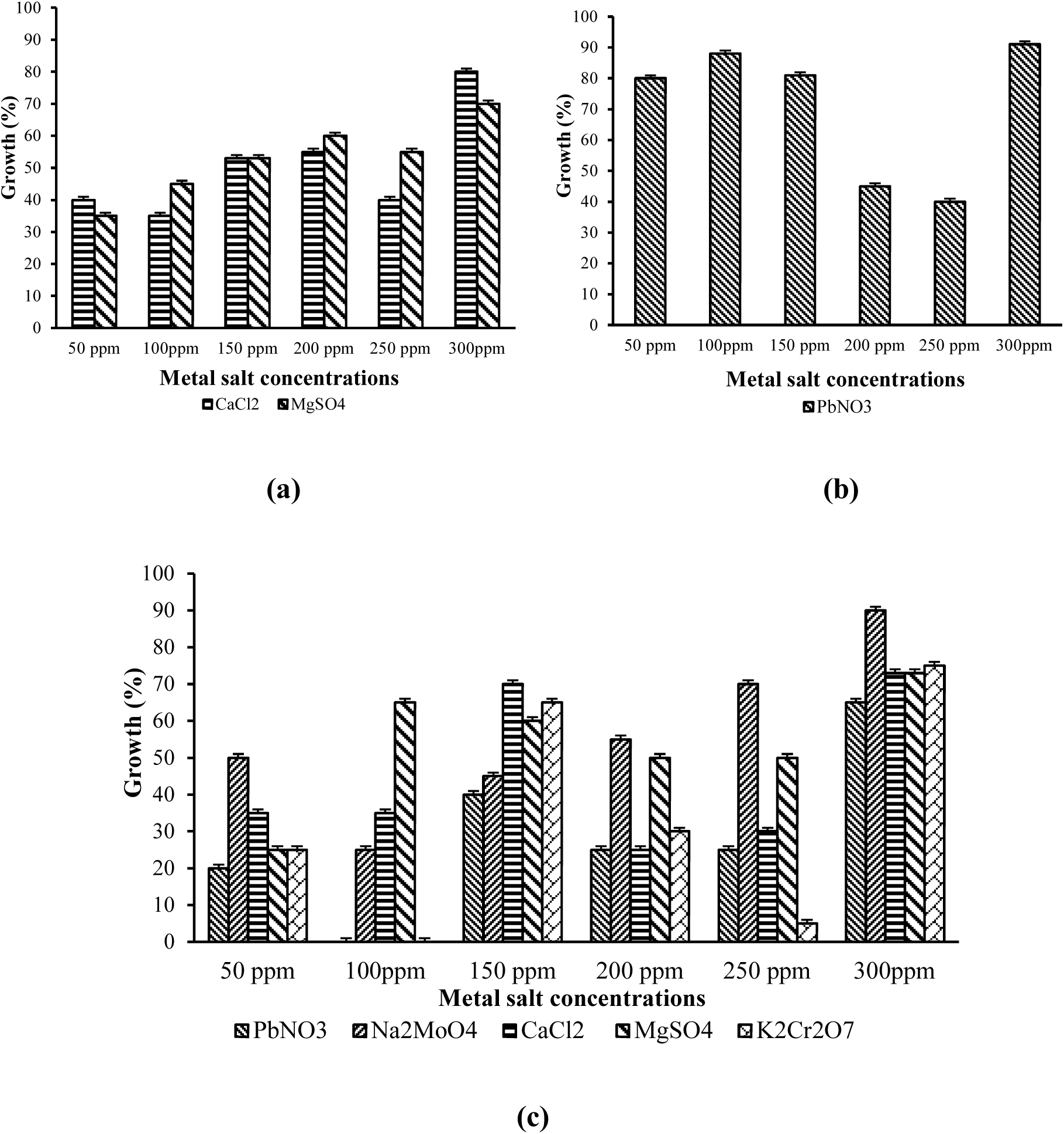
Maximum tolerance limits of bacterial isolates to metals salts solutions at 50, 100, 150, 200, 250 and 300 ppm concentrations (a) D6, (b) D7, (c) P1

### 3.5 Identification of the bacterial isolates

BLAST analysis indicated that strain D6 was a *Bacillus* species with 100% homology to *Bacillus paramycoides* (Table 3). Phylogenetic analysis reveals that it closely resembled *Bacillus pseudomycoides* (Figure 7) but formed a separate outgroup, indicating that the isolated species was phylogenetically distinct from the BLAST reference sequences. It was one of the nine novel species of the *Bacillus cereus* group reported by Liu et al. (2017). BLAST analysis indicated that strain D7 was also a *Bacillus* species with 99.86% homology to *Bacillus paramycoides* (Table 4). Phylogenetic analysis reveals that it closely resembled *Bacillus pseudomycoides* (Figure 8) but formed a separate outgroup, indicating that the isolated species was phylogenetically distinct from the BLAST sequences. Thus, D6 and D7 isolates share a high similarity. BLAST analysis indicated that strain P1 was an *Alcaligenes* species with 97.47% homology to *Alcaligenes faecalis* (Table 5). Phylogenetic analysis reveals that it closely resembled *Paenalcaligenes suuwonensis* and *Paenalcaligenes hominis* (Figure 9) but formed a separate outgroup, indicating that the isolated species was phylogenetically distinct from the BLAST sequences. The nucleotide sequences of these isolates D6, D7 and P1 have been submitted to GenBank under accession number [GenBank: MT477810], [GenBank: MT477812], and [GenBank: MT477813], respectively. The three isolates were then phylogenetically compared with each other and the top BLAST hit sequences. This result indicated that these isolates were more closely related to each other than the blast sequences. D6 and D7 clustered together demonstrating that these isolates were highly similar. The closest cluster was identified as *Paenalcaligenes suuwonensis* and *Paenalcaligenes hominis* (Figure 10).

**Table 3.** Top 10 BLAST hits for isolate D6.

| Hit Number | Description | Max Score | Total Score | Query Cover | Percentage Identity | Accession number |
| --- | --- | --- | --- | --- | --- | --- |
| 1. | *Bacillus paramycoides*<br>strain MCCC 1A04098 16S<br>ribosomal RNA, partial sequence | 2743 | 2743 | 100% | 100.00% | NR_157734.1 |
| 2. | *Bacillus tropicus*<br>strain MCCC 1A01406 16S<br>ribosomal RNA, partial sequence | 2737 | 2737 | 100% | 99.93% | NR_157736.1 |
| 3. | *Bacillus nitratireducens*<br>strain MCCC 1A00732 16S<br>ribosomal RNA, partial sequence | 2737 | 2737 | 100% | 99.93% | NR_157732.1 |
| 4. | *Bacillus luti*<br>strain MCCC 1A00359 16S<br>ribosomal RNA, partial sequence | 2737 | 2737 | 100% | 99.93% | NR_157730.1 |
| 5. | *Bacillus albus*<br>strain MCCC 1A02146 16S<br>ribosomal RNA, partial sequence | 2737 | 2737 | 100% | 99.93% | NR_157729.1 |
| 6. | *Bacillus cereus*<br>strain CCM 2010 16S ribosomal<br>RNA, partial sequence | 2732 | 2732 | 100% | 99.87% | NR_115714.1 |
| 7. | *Bacillus cereus*<br>ATCC 14579 16S ribosomal RNA<br>(rrnA), partial sequence | 2732 | 2732 | 100% | 99.87% | NR_074540.1 |
| 8. | *Bacillus proteolyticus*<br>strain MCCC 1A00365 16S<br>ribosomal RNA, partial sequence | 2726 | 2726 | 100% | 99.80% | NR_157735.1 |
| 9. | *Bacillus cereus*<br>strain IAM 12605 16S ribosomal<br>RNA, partial sequence | 2721 | 2721 | 99% | 99.86% | NR_115526.1 |
| 10. | *Bacillus wiedmannii*<br>strain FSL W8-0169 16S<br>ribosomal RNA, partial sequence | 2721 | 2721 | 100% | 99.73% | NR_152692.1 |

**Table 4.**
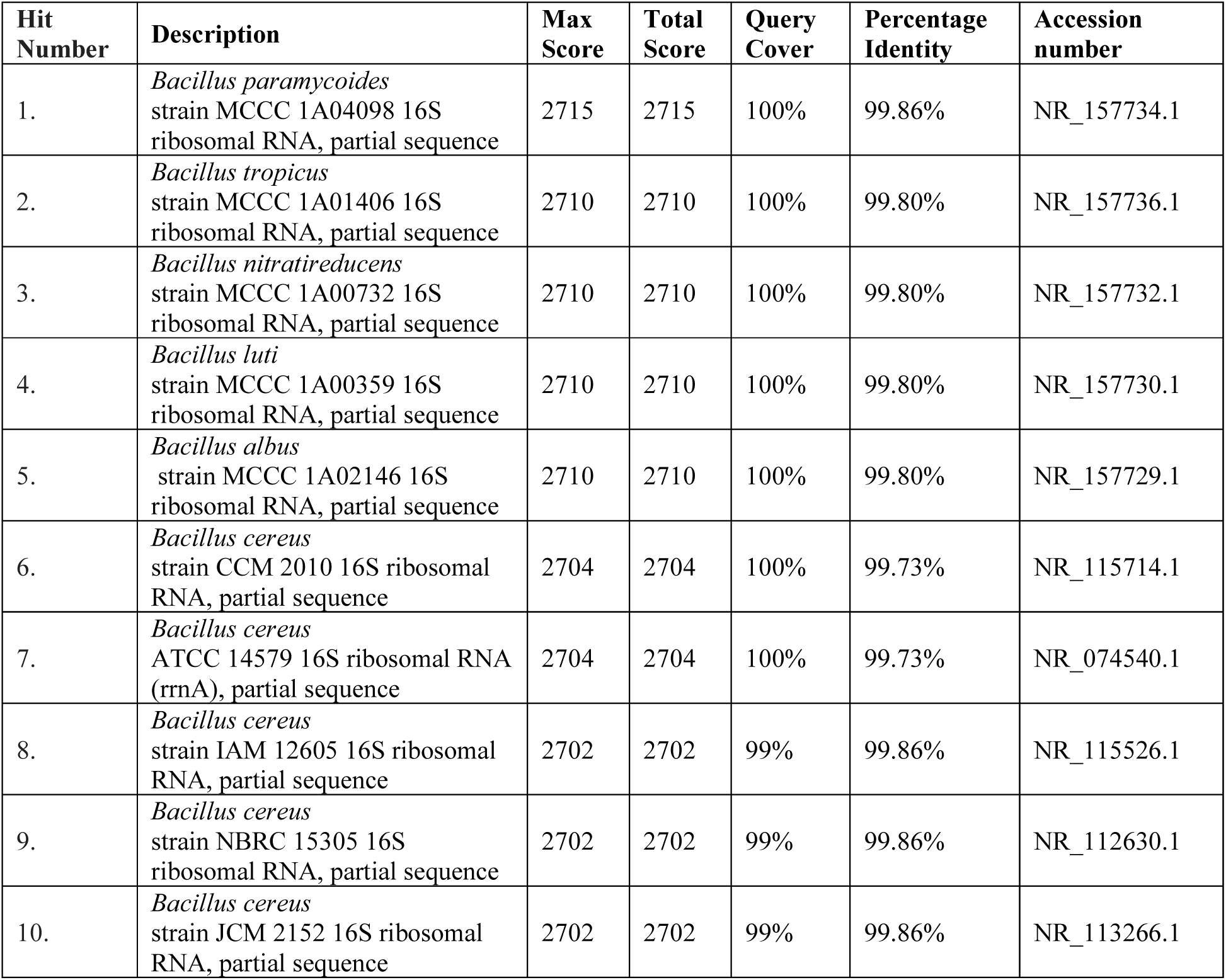
Top 10 BLAST hits for isolate D7.

**Table 5.**
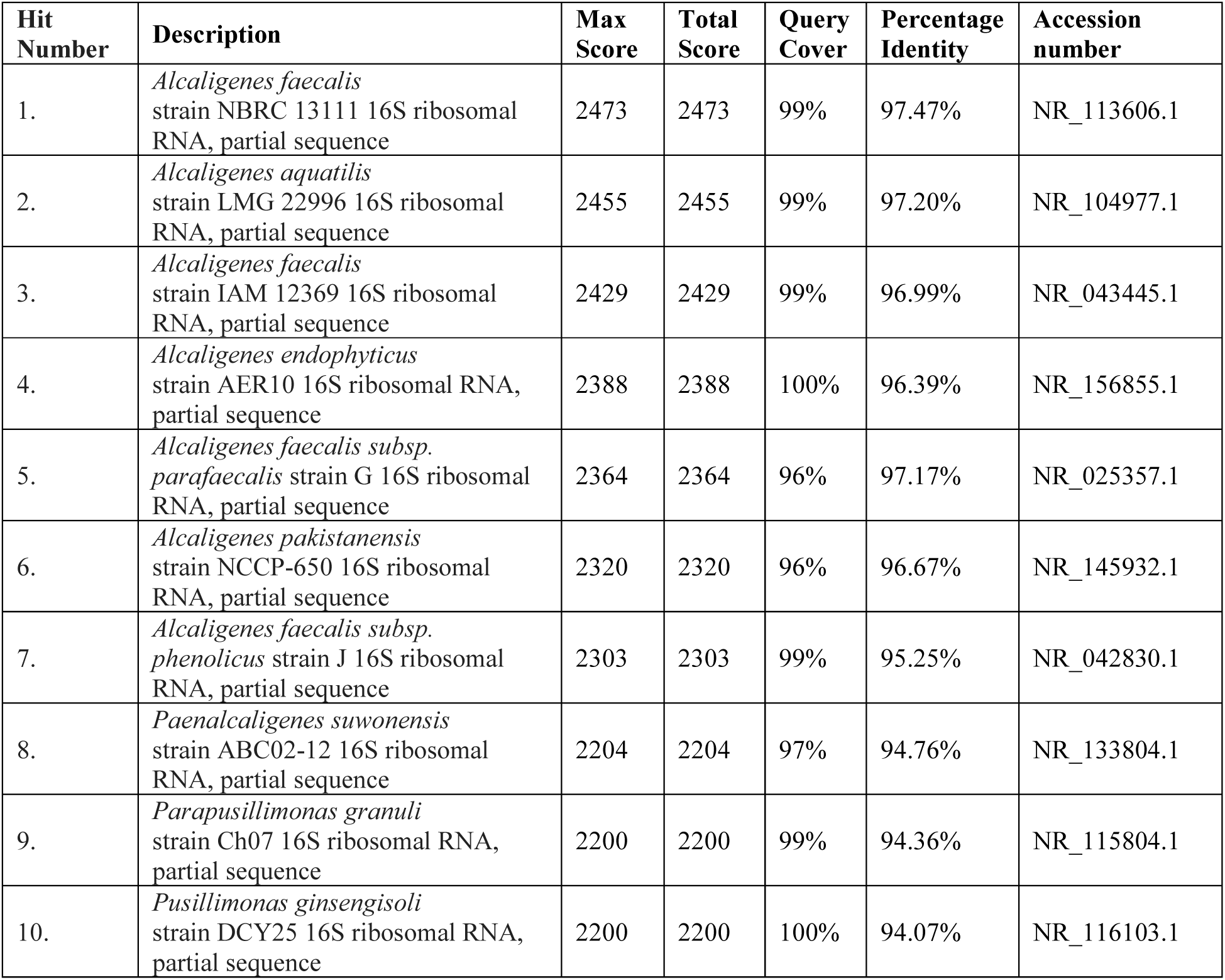
P1 Isolate Top 10 Blast hits.

| Hit Number | Description | Max Score | Total Score | Query Cover | Percentage Identity | Accession number |
| --- | --- | --- | --- | --- | --- | --- |
| 1. | *Alcaligenes faecalis* strain NBRC 13111 16S ribosomal RNA, partial sequence | 2473 | 2473 | 99% | 97.47% | NR_113606.1 |
| 2. | *Alcaligenes aquatilis* strain LMG 22996 16S ribosomal RNA, partial sequence | 2455 | 2455 | 99% | 97.20% | NR_104977.1 |
| 3. | *Alcaligenes faecalis* strain IAM 12369 16S ribosomal RNA, partial sequence | 2429 | 2429 | 99% | 96.99% | NR_043445.1 |
| 4. | *Alcaligenes endophyticus* strain AER10 16S ribosomal RNA, partial sequence | 2388 | 2388 | 100% | 96.39% | NR_156855.1 |
| 5. | *Alcaligenes faecalis* subsp. *parafaecalis* strain G 16S ribosomal RNA, partial sequence | 2364 | 2364 | 96% | 97.17% | NR_025357.1 |
| 6. | *Alcaligenes pakistanensis* strain NCCP-650 16S ribosomal RNA, partial sequence | 2320 | 2320 | 96% | 96.67% | NR_145932.1 |
| 7. | *Alcaligenes faecalis* subsp. *phenolicus* strain J 16S ribosomal RNA, partial sequence | 2303 | 2303 | 99% | 95.25% | NR_042830.1 |
| 8. | *Paenalcaligenes suwonensis* strain ABC02-12 16S ribosomal RNA, partial sequence | 2204 | 2204 | 97% | 94.76% | NR_133804.1 |
| 9. | *Parapusillimonas granuli* strain Ch07 16S ribosomal RNA, partial sequence | 2200 | 2200 | 99% | 94.36% | NR_115804.1 |
| 10. | *Pusillimonas ginsengisoli* strain DCY25 16S ribosomal RNA, partial sequence | 2200 | 2200 | 100% | 94.07% | NR_116103.1 |

**Figure 7.**
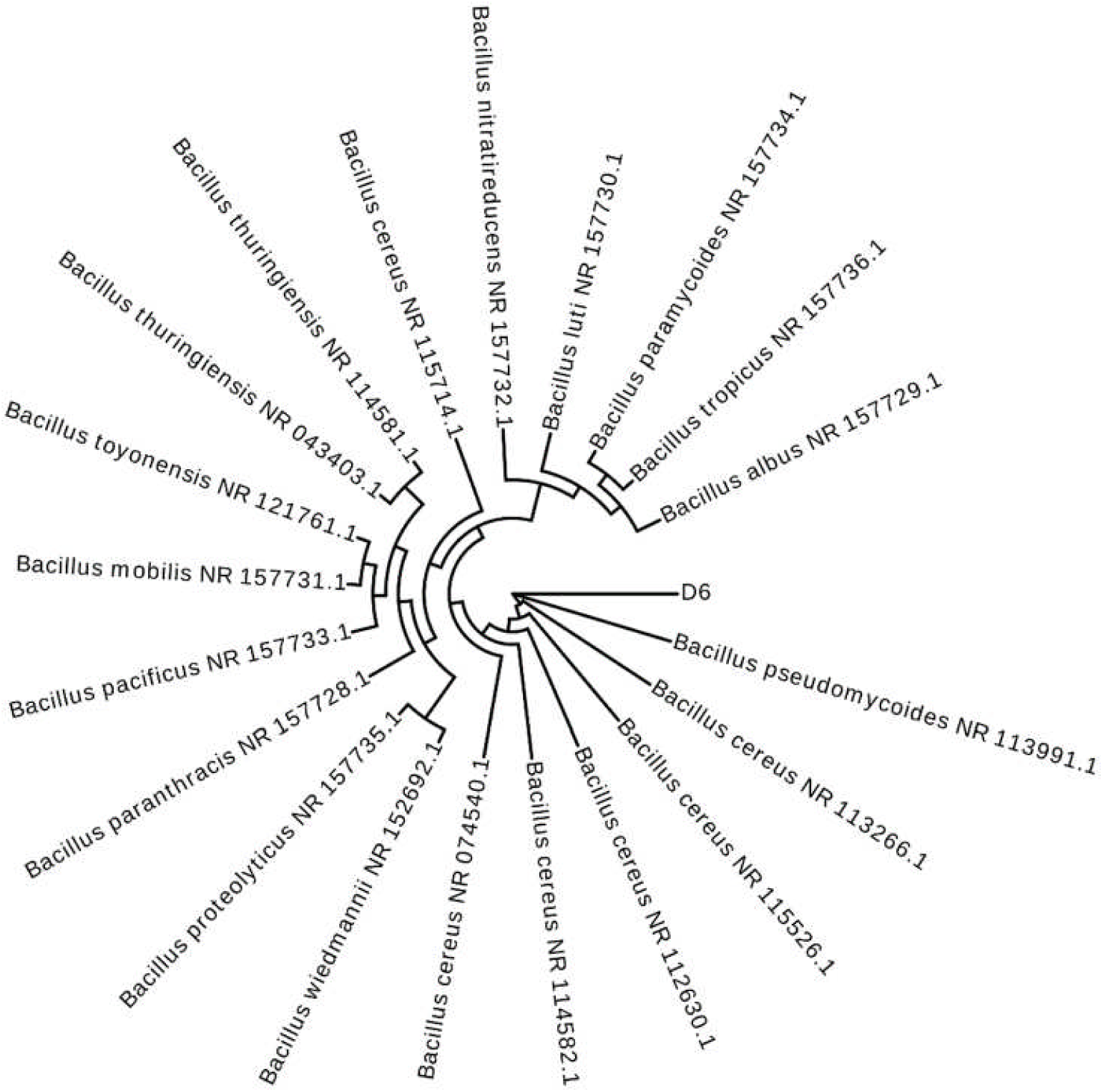
Phylogenetic distance between D6 isolate and top 20 BLAST sequences.

**Figure 8.**
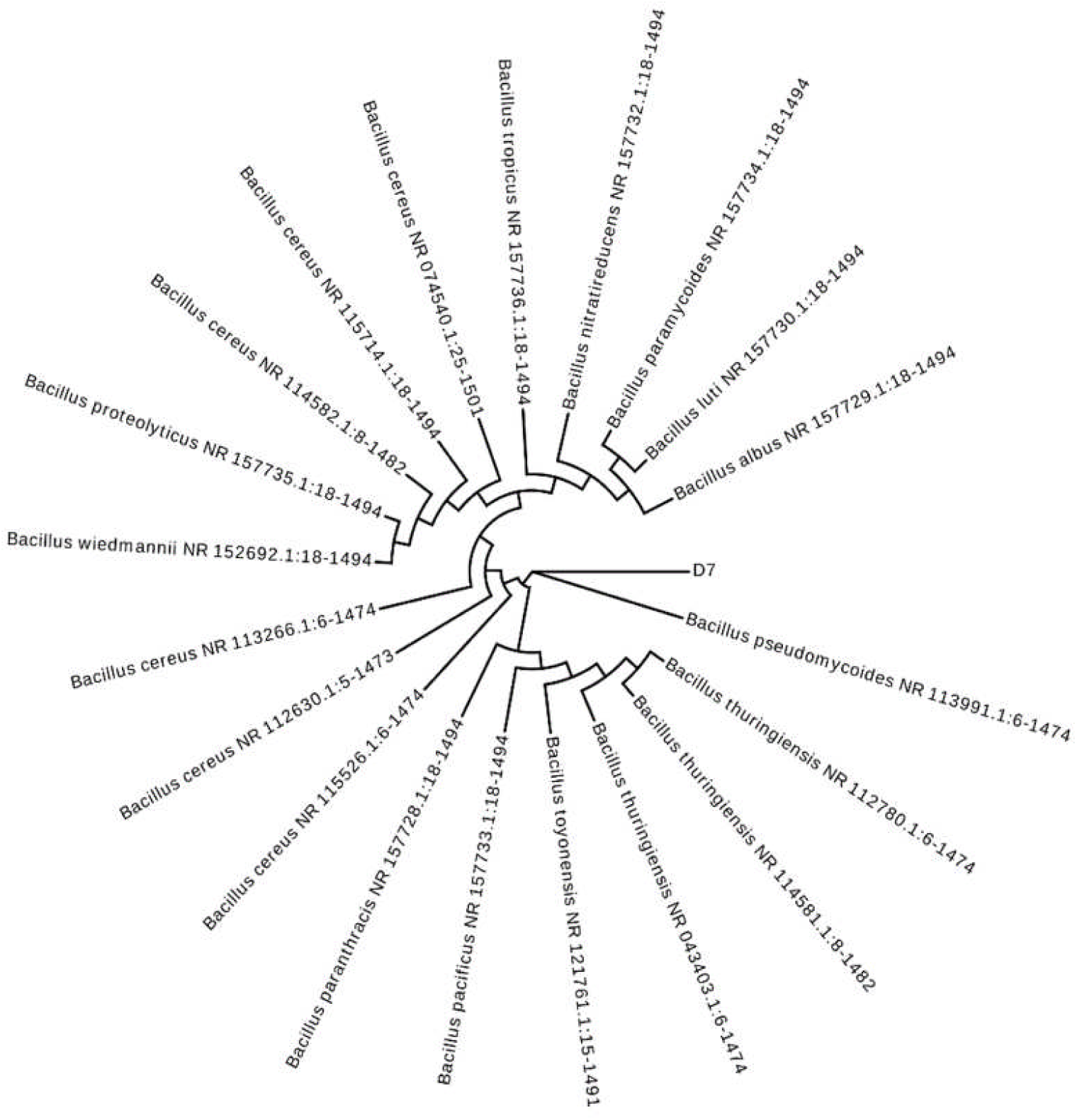
Phylogenetic distance between D7 isolate and top 20 BLAST sequences.

**Figure 9.**
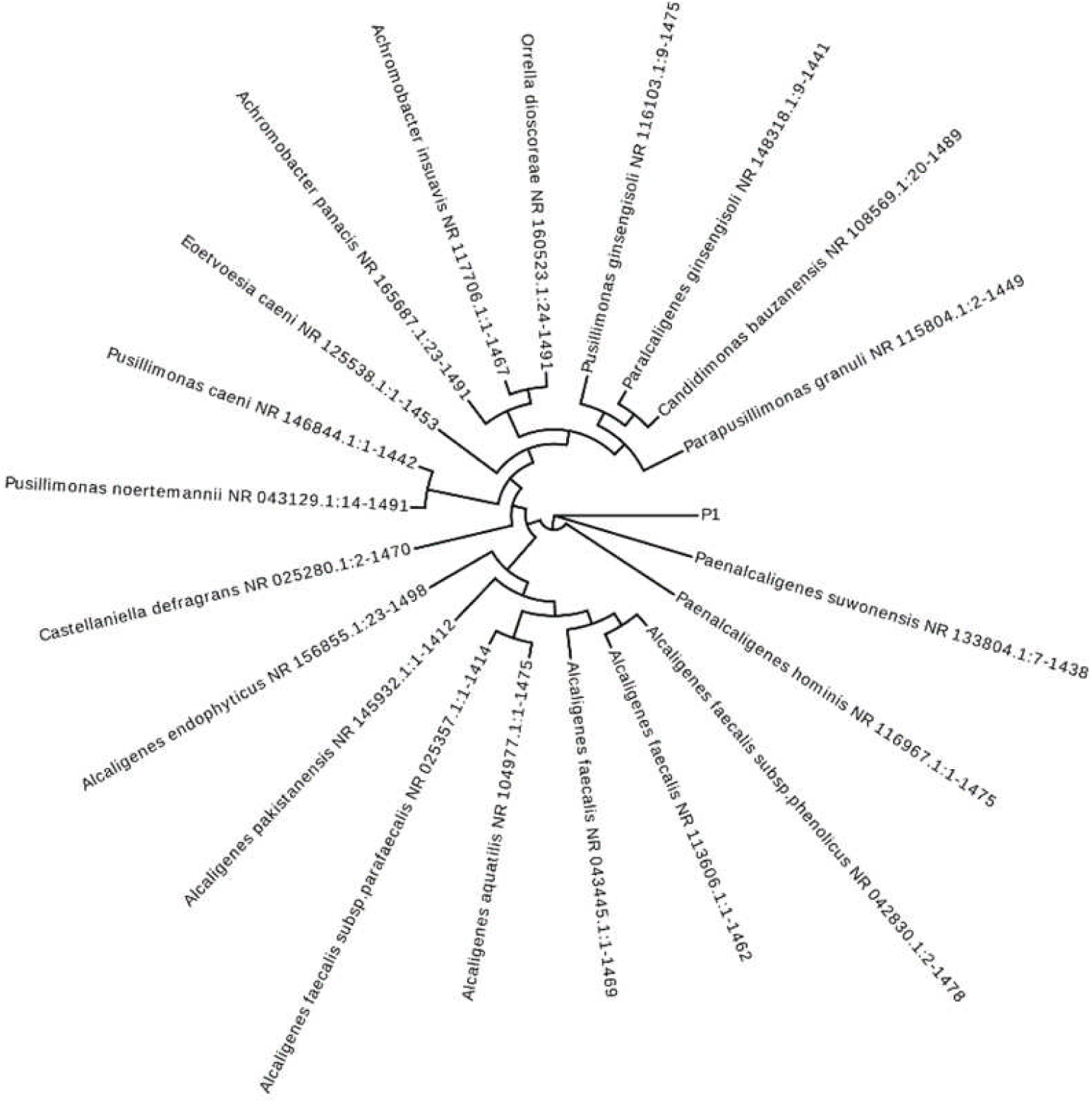
Phylogenetic distance between P1 isolate and top 20 BLAST sequences.

**Figure 10.**
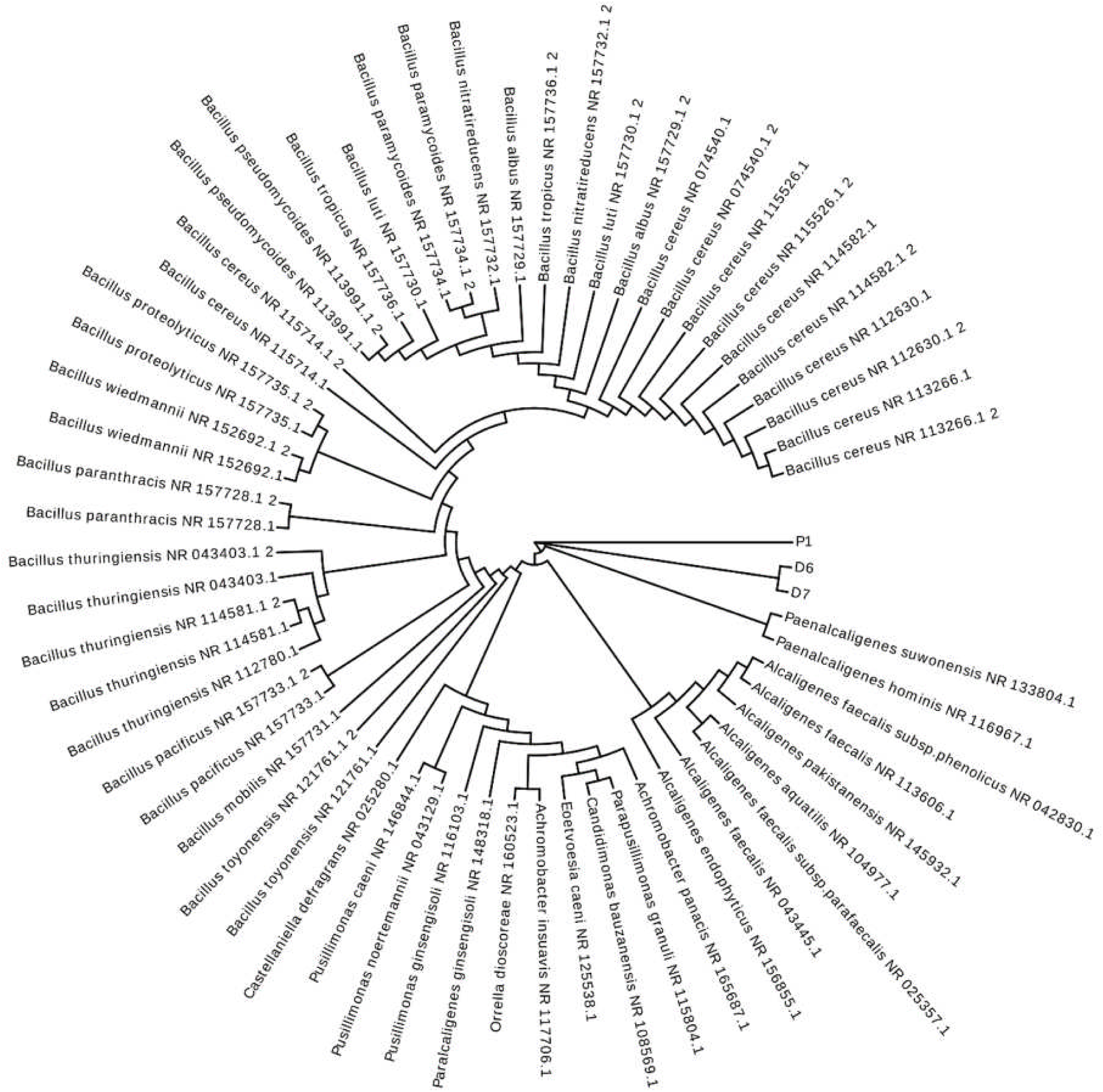
Phylogenetic relationship between the three isolates (D6, D7, P1) and BLAST reference sequences.

Authors have previously found *Bacillus* species such as *Bacillus paramycoides* to be part of plant growth-promoting rhizobacteria (Osman and Yin, 2018) and associated with bioremediation of toxic effluents containing cyanide (Wu et al., 2014), alkylphenols (Chang et al., 2020) and hydrocarbons (Kostka et al., 2011). Similarly, *Alcaligenes faecalis* was also noted as a biocontrol agent by Yokoyama et al. (2013). It must be noted that both species are potential human pathogens (Bottone, 2010; Kaliaperumal et al., 2006). The potential pathogenicity of these isolates would warrant further investigation prior to any bioamendment strategies.

It is very important to highlight that the optimal incubation time for *B. paramycoides* has not been reported previously. For *A. faecalis* JBW4, isolated from activated sludge, the optimal incubation time was 5 days (Kong et al., 2013). In present study, these two strains have proven to be very efficient in terms of requiring less time of incubation (48 hours) with more decolourisation potential. The optimal temperature for growth of *Bacillus* spp. is reported between 30 – 37 °C (Gilbert et al., 2009). *Alcaligenes faecalis* was previously reported to be grown at 37 °C (Schroll et al., 2001). Syed et al. (2015) have found heavy metal resistance and their degradation by *B. cereus* strains. *A. faecalis* was found to be heavy metal resistant bacteria isolated from sewage wastewater and responsible for the synthesis of silver nanoparticles (Abo-Amer et al., 2015). The capability of *A. faecalis* to degrade phenol as a carbon source has been previously reported (Rehfuss and Urban 2005). This supports our results showing biodegradation of phenol into other non-toxic low molecular organic compounds. *A. faecalis* has been proven to be efficient to bioremediate ε-Caprolactam too from nylon-6 produced wastewater plant (Baxi and Shah 2002). But to the best of the authors knowledge, *B. paramycoides* have never been reported for any type of wastewater bioremediation. The antibiotic degradation potential of different isolated bacterial species from pharmaceutical wastewaters (Tahrani et al., 2015) and the biodegradation of acrylamide by *Enterobacter aerogenes* isolated from domestic wastewater (Buranasilp and Charoenpanich, 2011) has only been reported previously. Majorly, they looked at the individual wastewaters while our work has investigated a complex combination and is more representative of the real-world scenario in Pakistan.

## 4. Environmental implications

Our work suggests that *B. paramycoides* D6, *B. paramycoides* D7 and *A. faecalis* are capable to bioremediate domestic, hospital, textile, pharmaceutical and mixed wastewaters under optimal conditions. These optimal conditions for temperature (37 and 51 °C) are achievable in Pakistan’s arid climate (in temperate zone) and the incubation time is achieved in 48 h only. The utilization of these bacterial strains has several advantages as compared to the conventional methods such as physicochemical approaches for the removal of contaminants. Bacterial treatment with these strains is a cost-effective and low-tech method as the strains are isolated from the same wastewater needed to be treated (Phugare et al., 2011). Further, these bacterial strains have been found here to be efficient for the biotreatment of a wide range of wastewaters, *i.e*. domestic, hospital, textile, pharmaceutical and mixed wastewaters. The strains degraded pharmaceutic pollutants into ecofriendly derivatives and showed high decolourisation potential. Thus, this work suggests that the biological treatment of wastewaters using *B. paramycoides* and *A. faecalis* can be an eco-friendly and efficient method which may help developing countries such as Pakistan to meet the Sustainable Development Goal of Clean Water and Sanitation (SDG-6). Future work may require to focus on scaling-up this methodology at commercial level and to form a consortium of these strains for achieving much higher efficiency.

## 5. Conclusion

Bacterial strains *B. paramycoides* D6, *B. paramycoides* D7 and *A. faecalis* have been proven to be efficient in terms of possessing bioremediation potential against different wastewaters, *i.e*. domestic, hospital, textile, pharmaceutical and mixed wastewaters. These bacterial isolates significantly biodegrade the pollutants from the wastewaters into non-toxic organic compounds within 48 hours of incubation, 10 % of inoculum and 37 and 51°C temperatures, respectively. Under these optimal growth conditions, the strains *B. paramycoides* D6, *B. paramycoides* D7 and *faecalis* showed maximum decolourisation potential of 96, 96, 93 %, respectively against hospital wastewater. GCMS analysis confirmed the biodegradation of pharmaceutic pollutants, *i.e*. Phenol, Salicylic acid, Caffeine, Naproxen, Octadecene and Diazepam, present in the hospital wastewater into Ticlopidine in the case of *B. paramycoides* D6, Tetradecene and Griseofulvin in the case of *B. paramycoides* D7 and Lidocaine and Butalbital in the case of *A. faecalis*. At 300 mg/L concentration, *B. paramycoides* D6, showed overall maximum growth of 78 and 70 % for CaCl_2_ and MgSO_4_, respectively; *B. paramycoides* D7 showed maximum growth of 82 % for PbNO_3_; *Alcaligenes faecalis* showed maximum growth of 65, 90, 73, 73 and 75 % for PbNO_3_, Na_2_MoO_4_, CaCl_2_, MgSO_4_ and K_2_Cr_2_O_7_, respectively. Our work recommends that the development of a consortium from these strains may prove more efficient source of bioremediation of wastewaters.

## Acknowledgements

Authors acknowledge the Department of Botany and GC University Lahore for their continuous support throughout the research work.

