## Supplementary Data for "Indigenous *Bacillus paramycoides* and *Alcaligenes faecalis*: potential solution for the bioremediation of wastewaters"

**Supplementary Information**

1. *Results*

*1.1 Optimal growth conditions*

The strain D6 showed the maximum decolourisation at 37 ℃ while lowest at 51℃. The same pattern was observed for D7 (Figure 1). The strains D6 and D7 were proven to be perfect mesophile in nature as they showed maximum decolourisation of 50 and 73 % at 37 ᵒC. The strain P1 showed maximum decolourisation of 78 % at 51 ℃. The present result indicated the presence of species at 51 °C and showed its existence as a thermophile. A big difference in decolourisation was observed between 51 and 58 degrees for all isolates even the thermophile. Perhaps some hydrolysis or production of toxic compounds may responsible for that.

Decolourisation (%)

| Temperature (℃) | Bacillus paramycoides D6 | Bacillus paramycoides D7 | Alcaligenes faecalis P1 |
| --- | --- | --- | --- |
| 30 | 23.6364 | 45.4545 | 70 |
| 37 | 69 | 80 | 51 |
| 44 | 9.09091 | 65.4545 | 60 |
| 51 | 1.3 | 30 | 90 |
| 58 | 38.1 | 56.3636 | 28 |

**Figure S1. Optimisation I: Temperature (℃)**

After 48 hours of incubation, the strains D6, D7 and P1 showed maximum decolourisation of 88, 88, 89 % against domestic wastewater, respectively (Figure 2).

Decolourisation (%)

| Time (h) | Bacillus paramycoides D6 | Bacillus paramycoides D7 | Alcaligenes faecalis P1 |
| --- | --- | --- | --- |
| 24 | 79.2553 | 86.4362 | 86.9681 |
| 48 | 87.766 | 88.2979 | 89.3617 |
| 72 | 72.6064 | 80.0532 | 81.383 |
| 96 | 81.9149 | 80.5851 | 86.1702 |

**Figure S2. Optimisation II: Incubation time (h)**

The bacterial cell count calculated for strains D6, D7 and P1 were 4.675×10^7^ cells/mL, 5.15×10^7^ cells/mL and 7.6×10^7^ cells/mL respectively. The strains D6, D7 and P1 showed maximum percentage decolourisation of 60, 68 and 70 for 10% of inoculum concentration (Figure 3). From previously reported work, Anto et al. (2006) and Shah et al. (2012) have reported the effectiveness of 10 % optimal inoculum size for the growth and decolourising activity of isolated bacterial strains.

Decolourisation (%)

| Inoculum concentraion (%) | Bacillus paramycoides D6 | Bacillus paramycoides D7 | Alcaligenes faecalis P1 |
| --- | --- | --- | --- |
| 5 | 30 | 50 | 57 |
| 10 | 60 | 68 | 70 |
| 15 | 30 | 61 | 28 |
| 20 | 23 | 35 | 45 |
| 25 | 18 | 45 | 15 |
| 30 | 30 | 35 | 20 |

**Figure S3. Optimisation III: Inoculum concentration (%)**

1.2 *Mass Spectra Analysis*

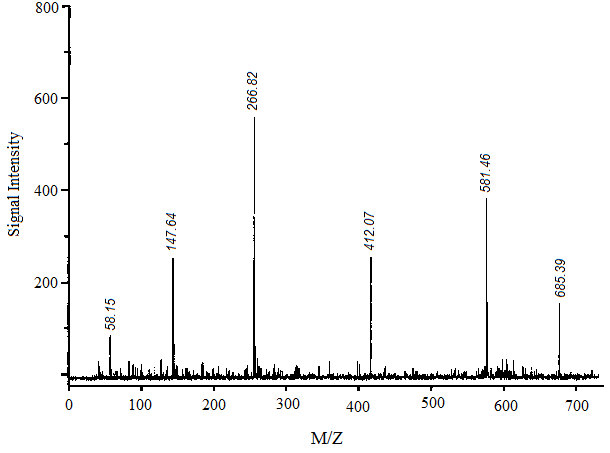

**Figure S4 (a). Mass spectrum of untreated hospital wastewater**

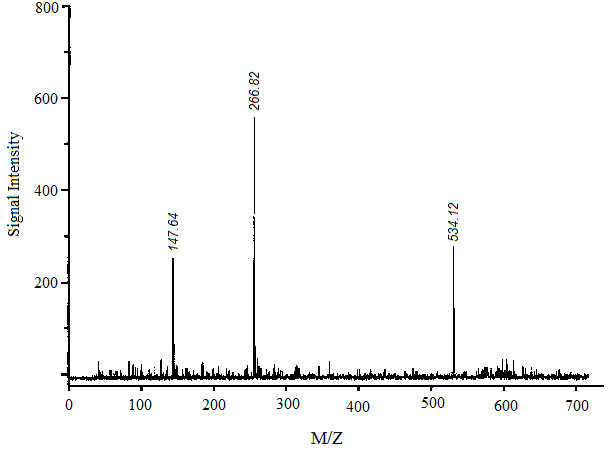

**Figure S4 (b). Mass spectrum of treated hospital wastewater with isolate D6**

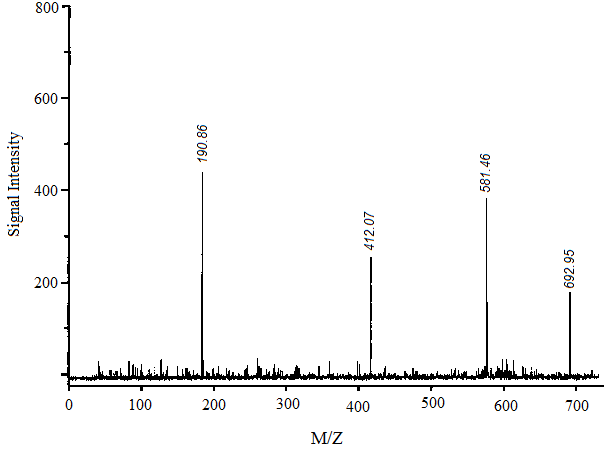

**Figure S4 (c). Mass spectrum of treated hospital wastewater with isolate D7**

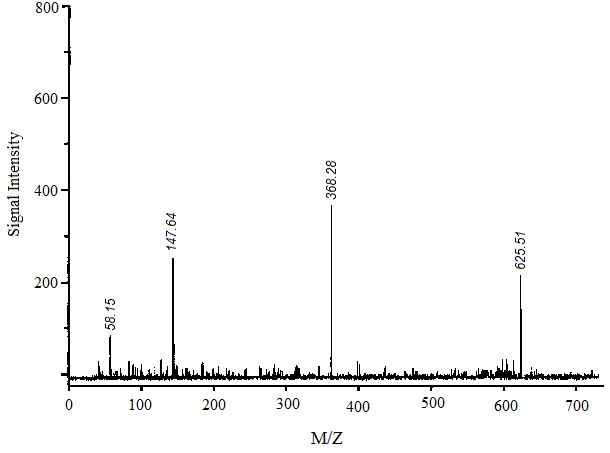

**Figure S4 (d). Mass spectrum of treated hospital wastewater with isolate P1**

*1.3 Identification of isolates*

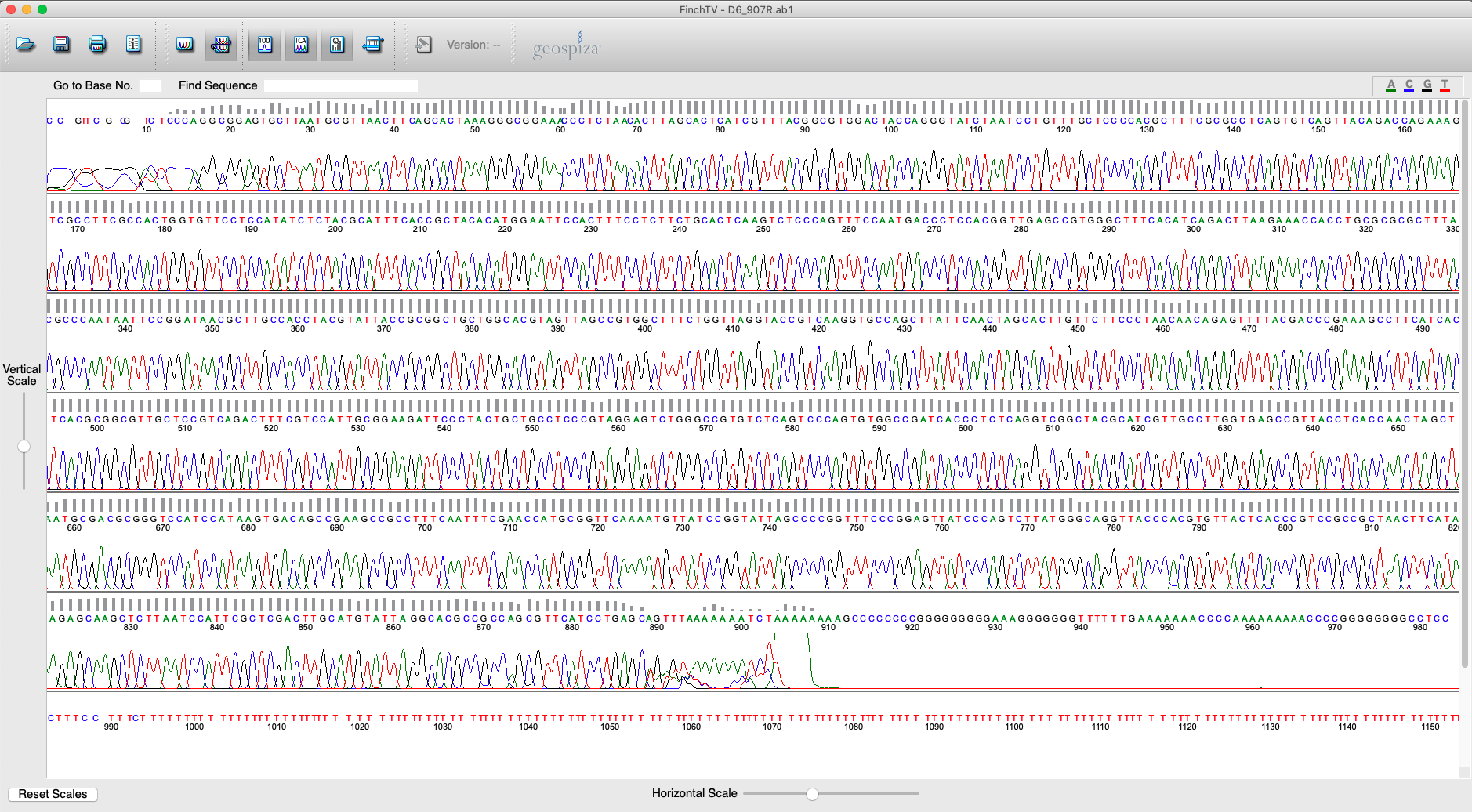

**Figure S5. D6 785F raw chromatogram**

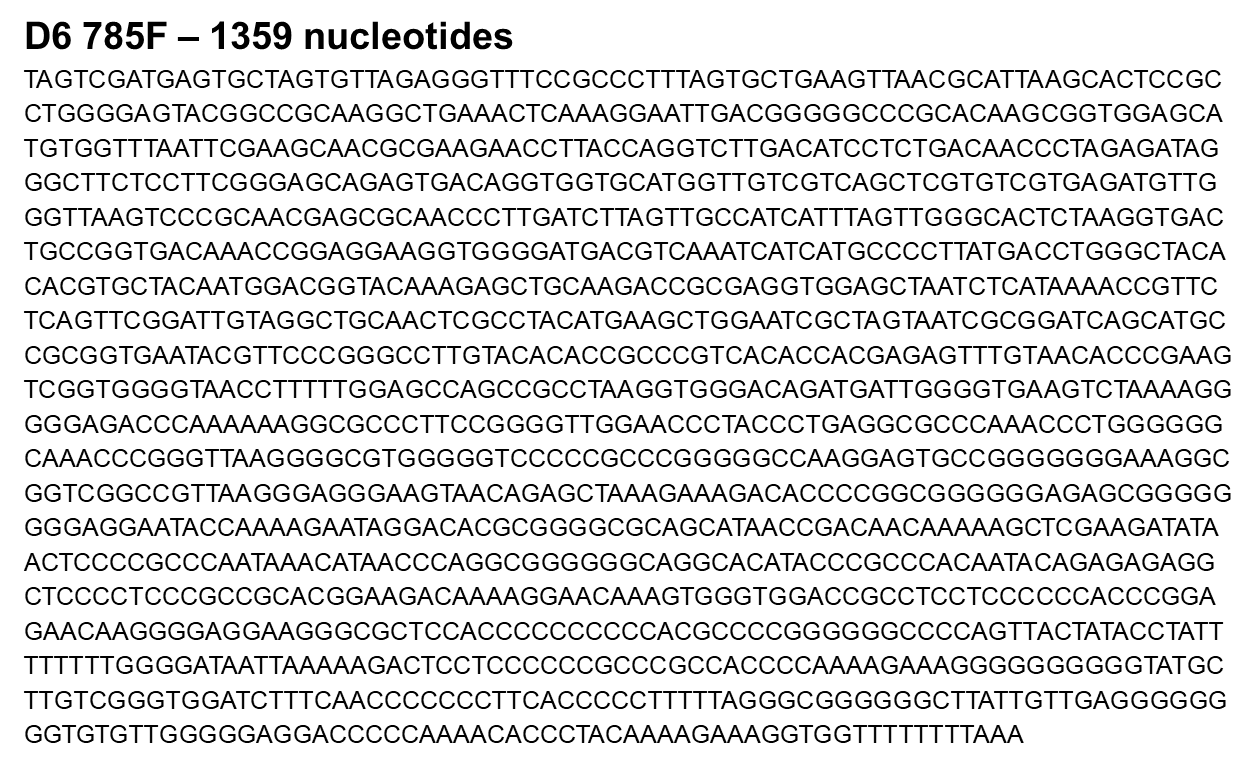

**Figure S6. D6 785F raw sequence**

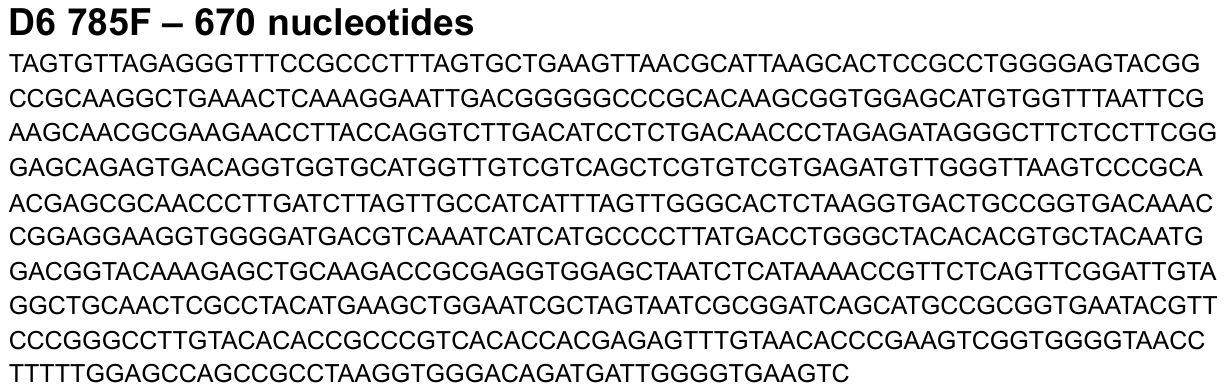

**Figure S7. D6 785F cleaned sequence**

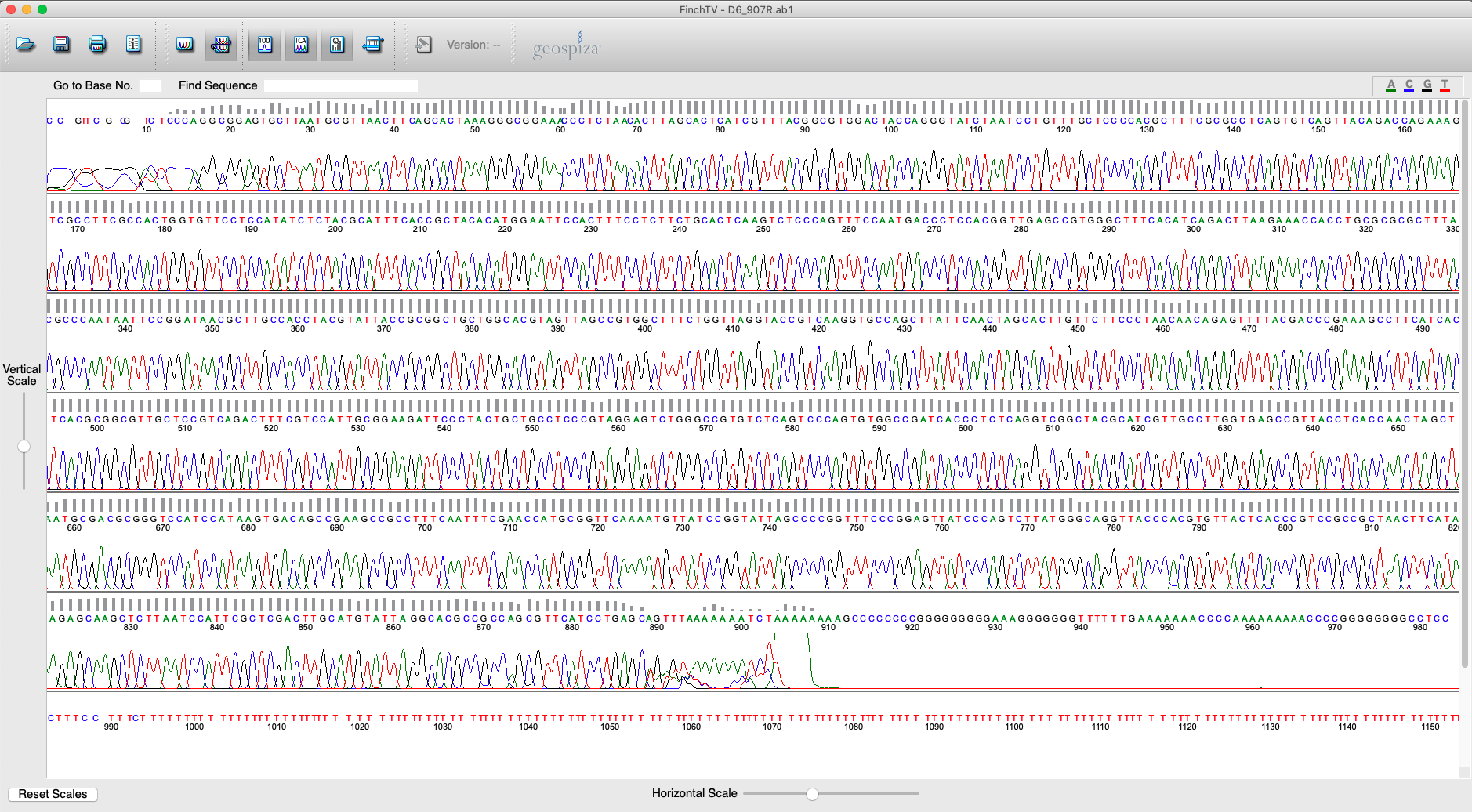

**Figure S8. D6 907R raw chromatogram**

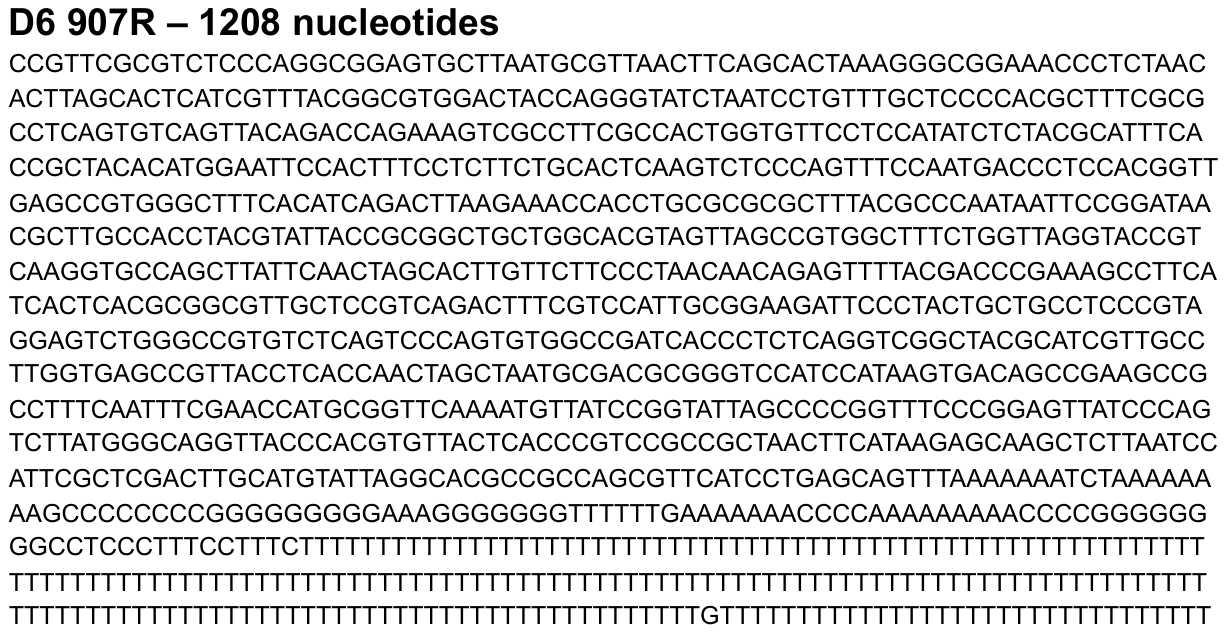

**Figure S9. D6 907R raw sequence**

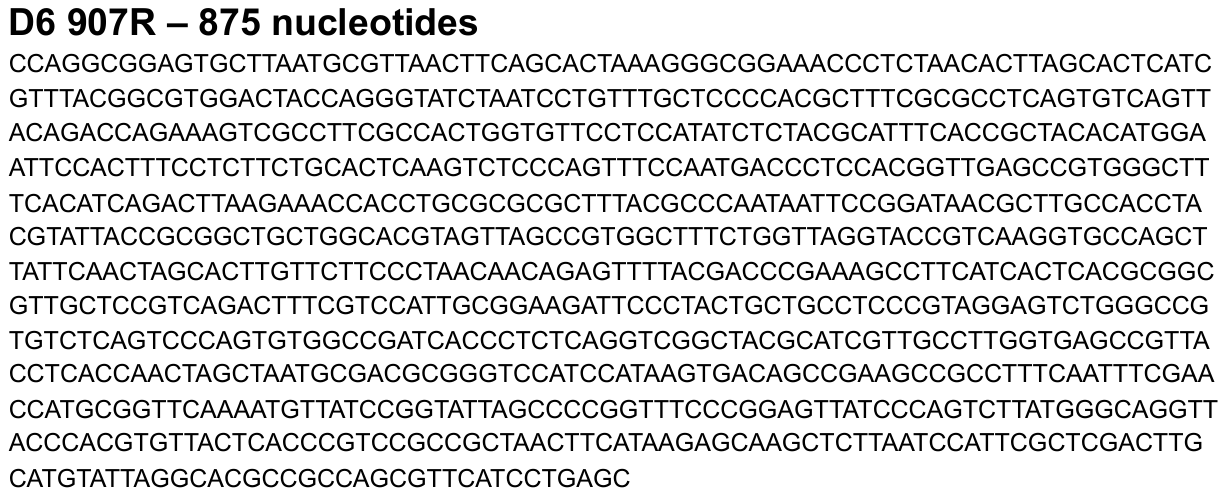

**Figure S10. D6 907R cleaned sequence**

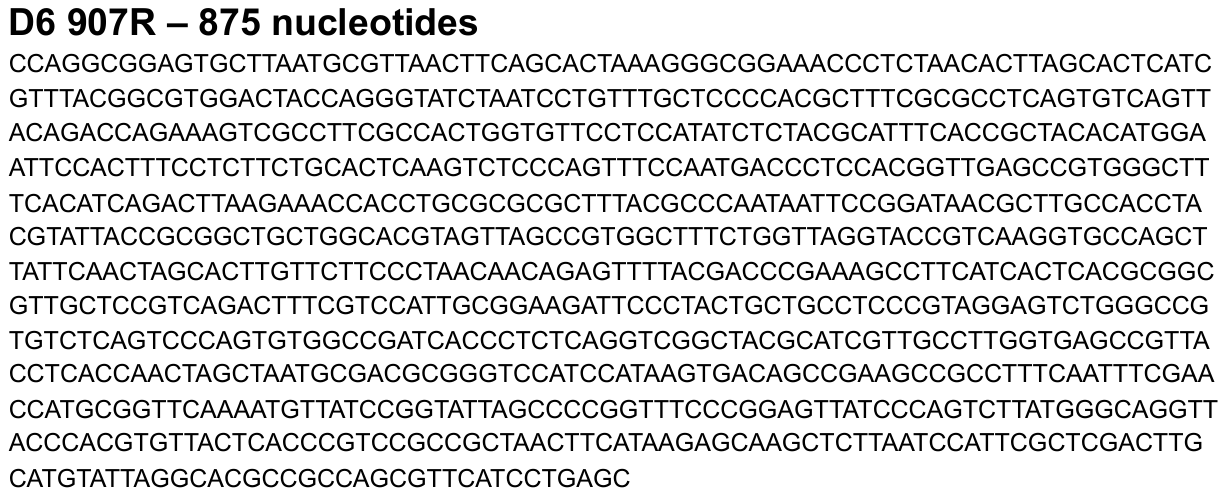

**Figure S11. D6 907R cleaned sequence**

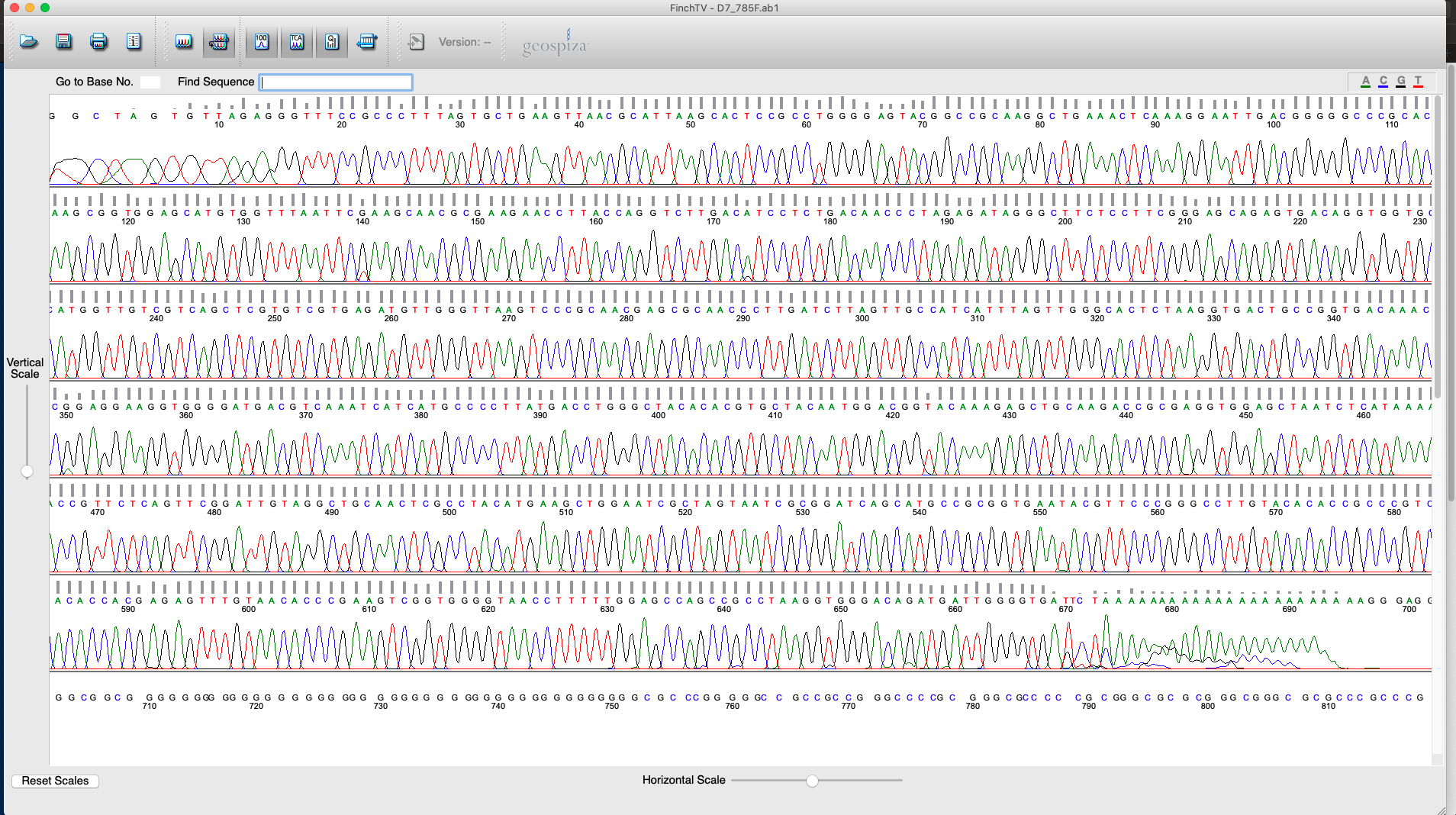

**Figure S12. D7 785F raw chromatogram**

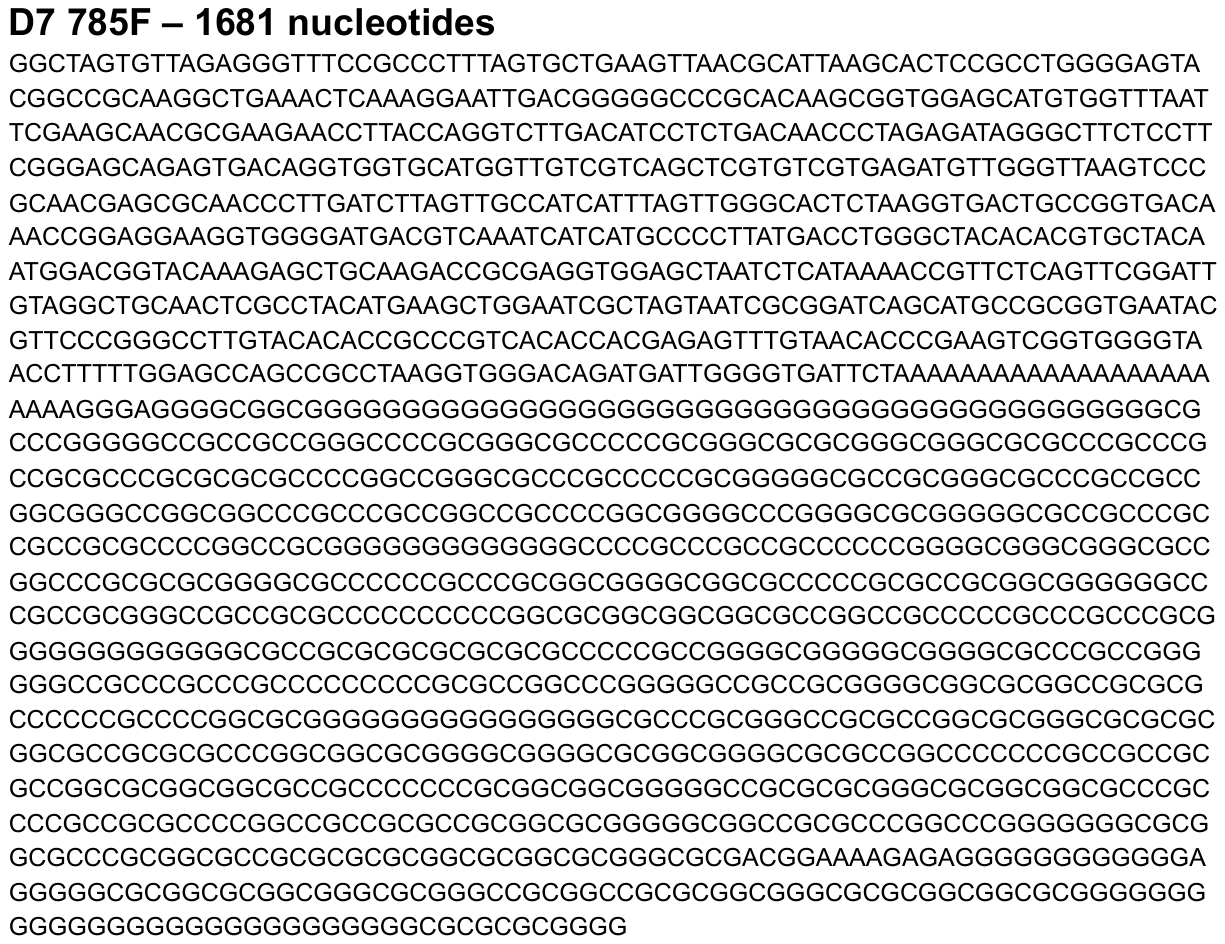

**Figure S13. D7 785F raw sequence**

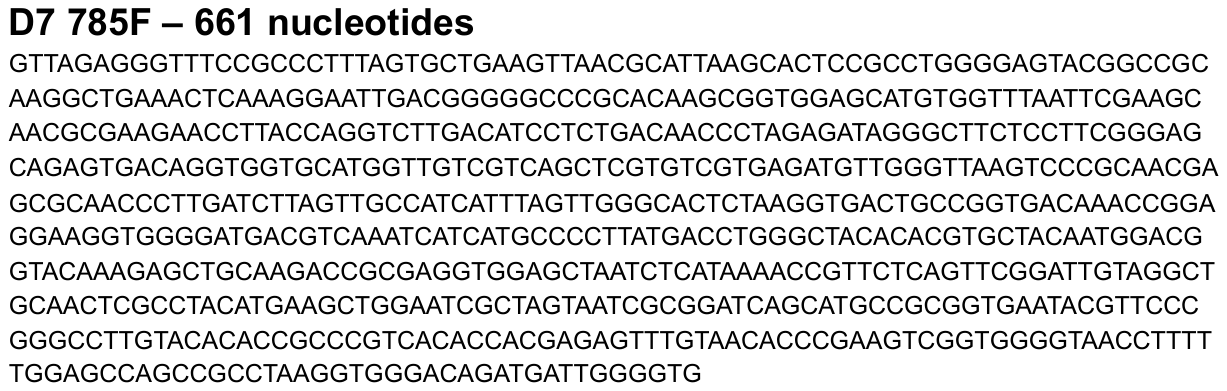

**Figure S14. D7 785F cleaned sequence**

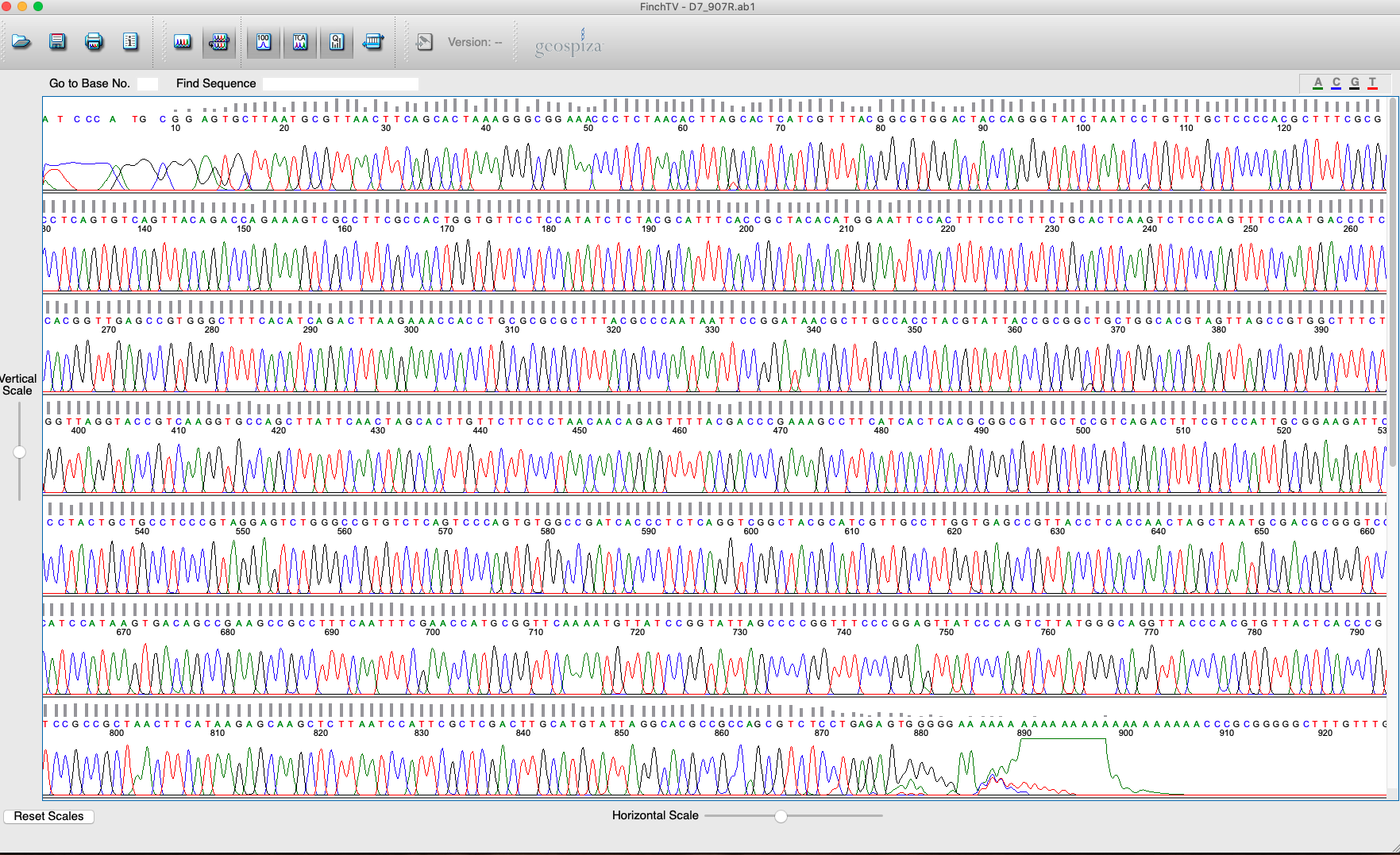

**Figure S15. D7 907R raw chromatogram**

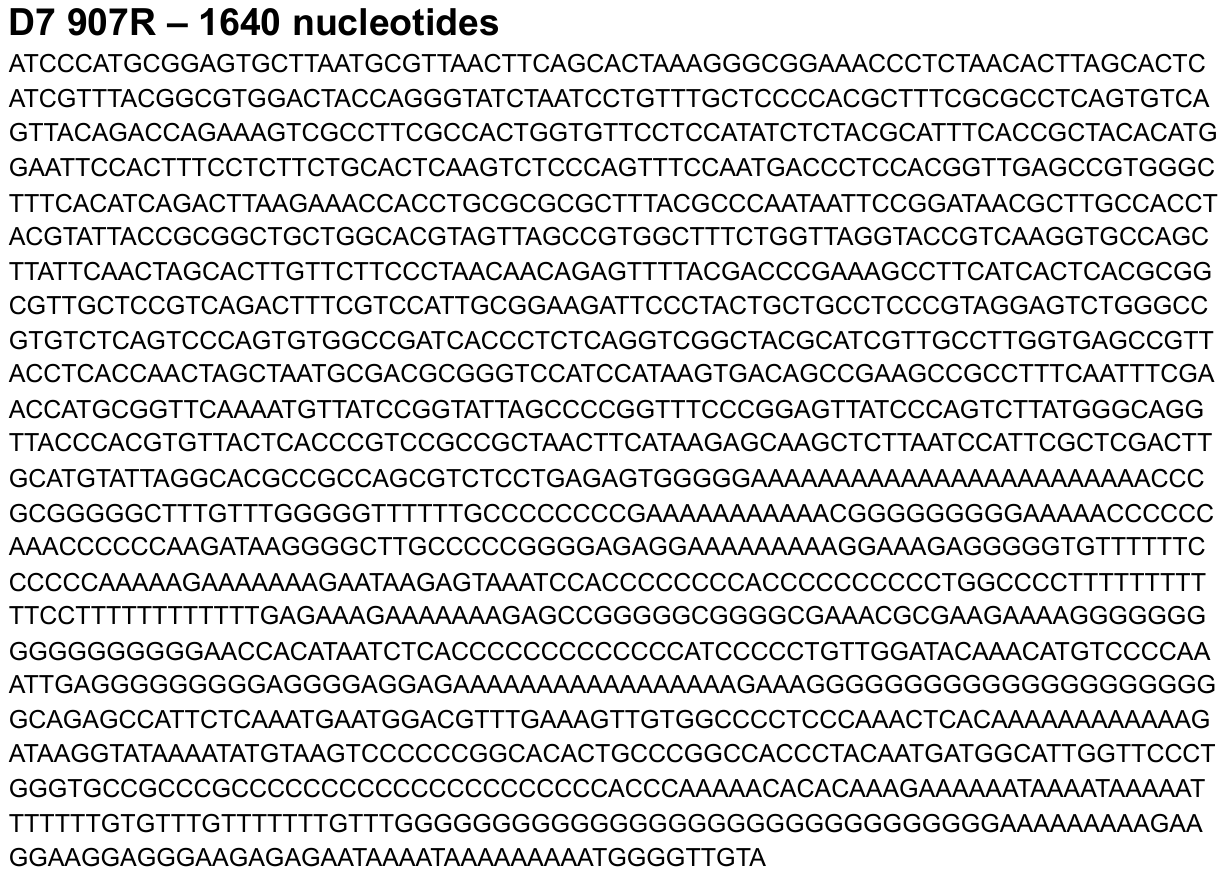

**Figure S16. D7 907R raw sequence**

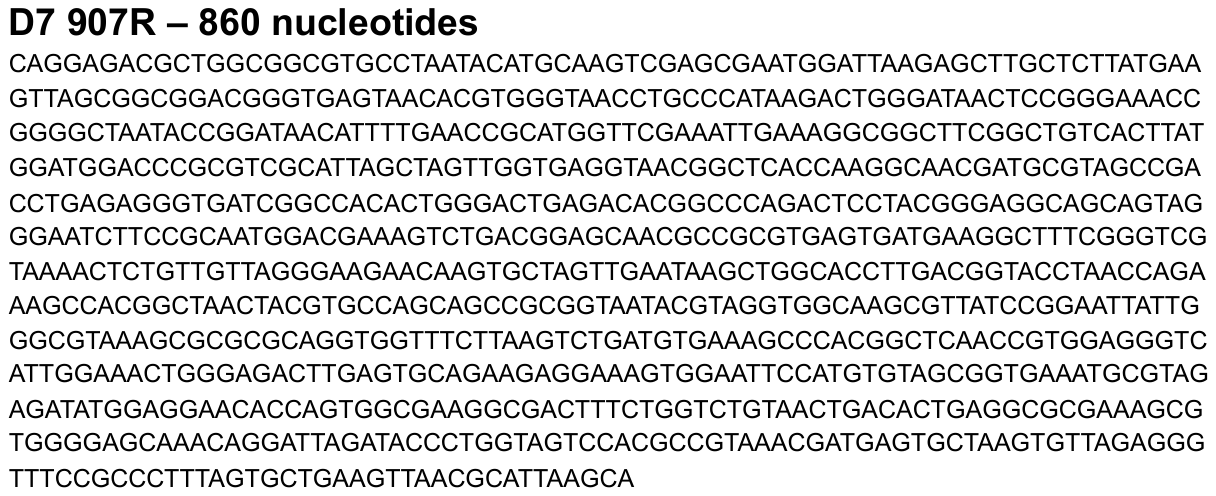

**Figure S17. D7 907R cleaned sequence**

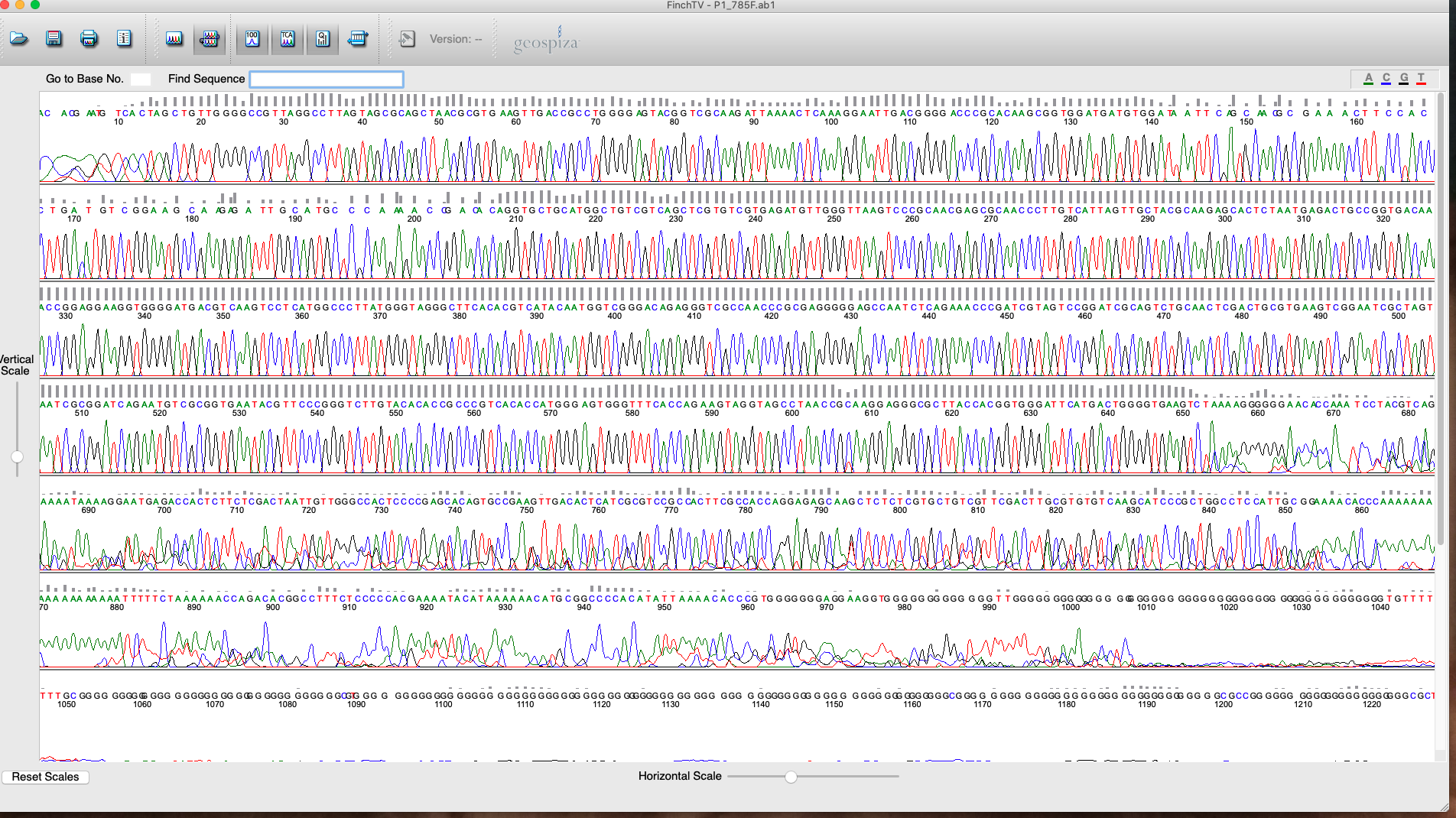

**Figure S18. P1 785F raw chromatogram**

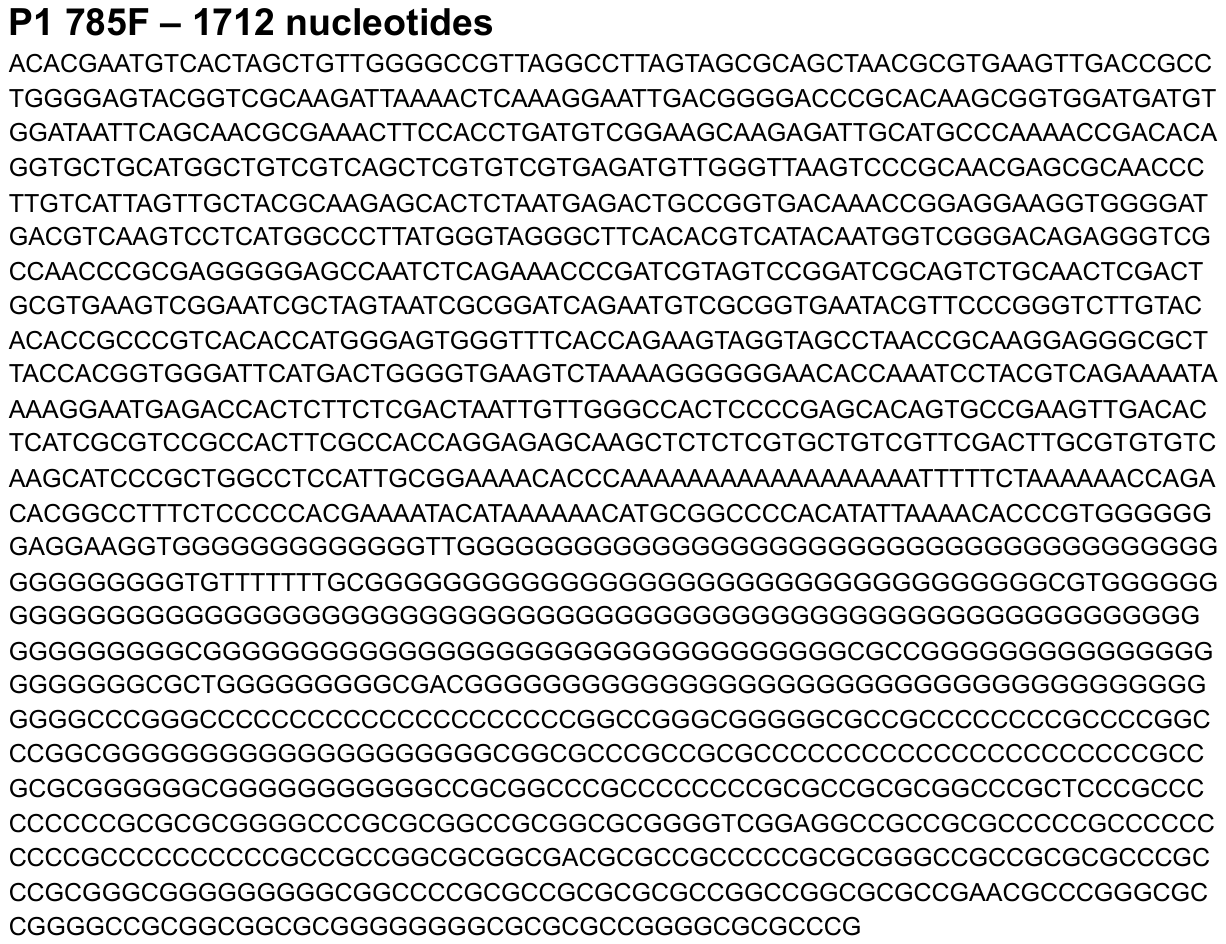

**Figure S19. P1 785F raw sequence**

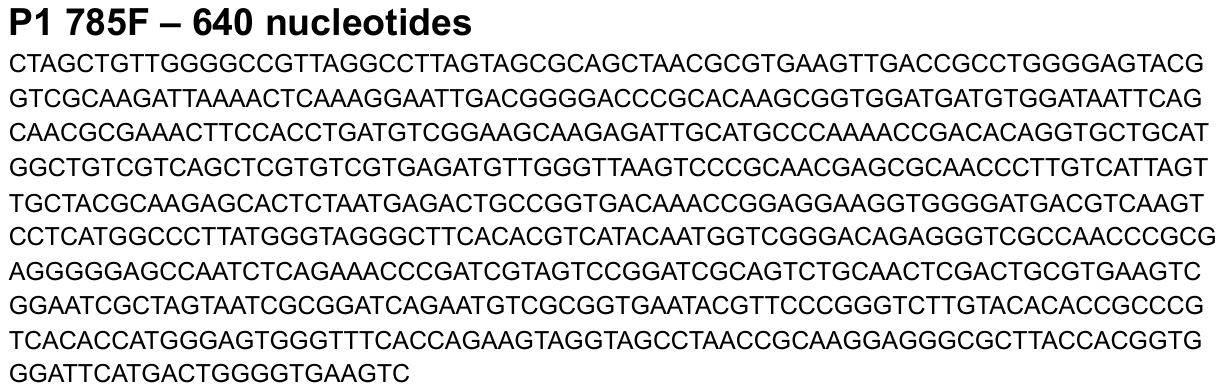

**Figure S20. P1 785F cleaned sequence**

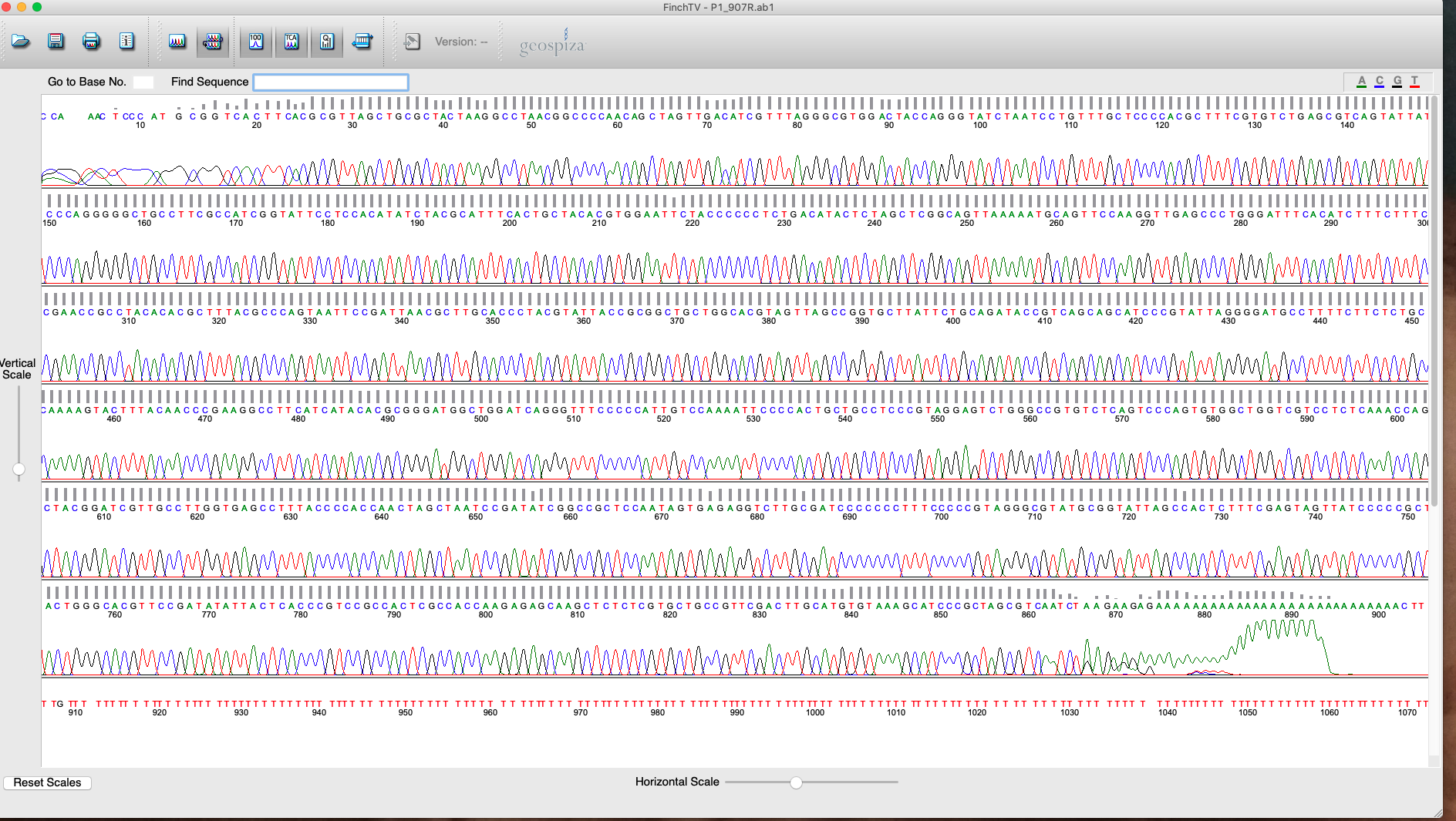

**Figure S21. P1 907R raw chromatogram**

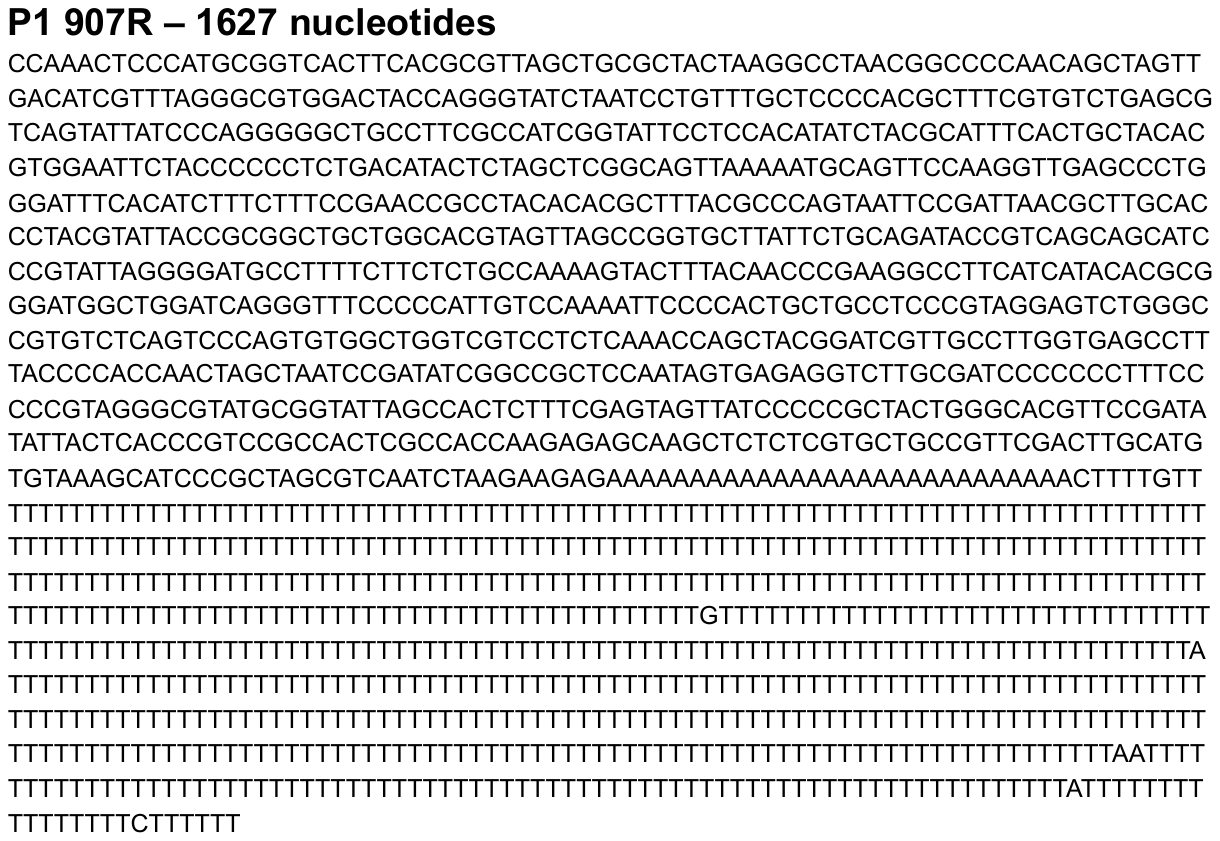

**Figure S22. P1 907R raw sequence**

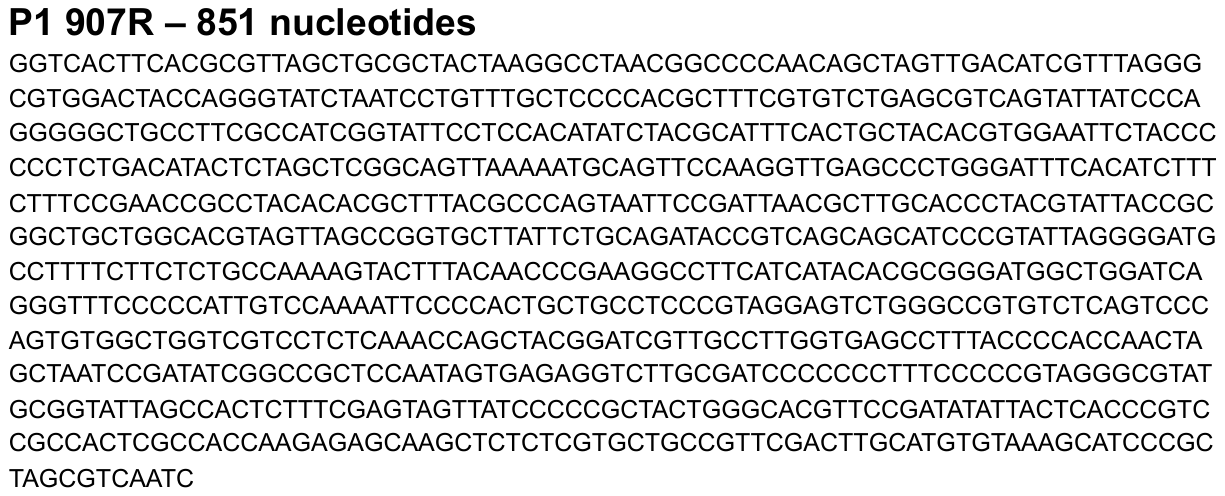

**Figure S23. P1 907 cleaned sequence**

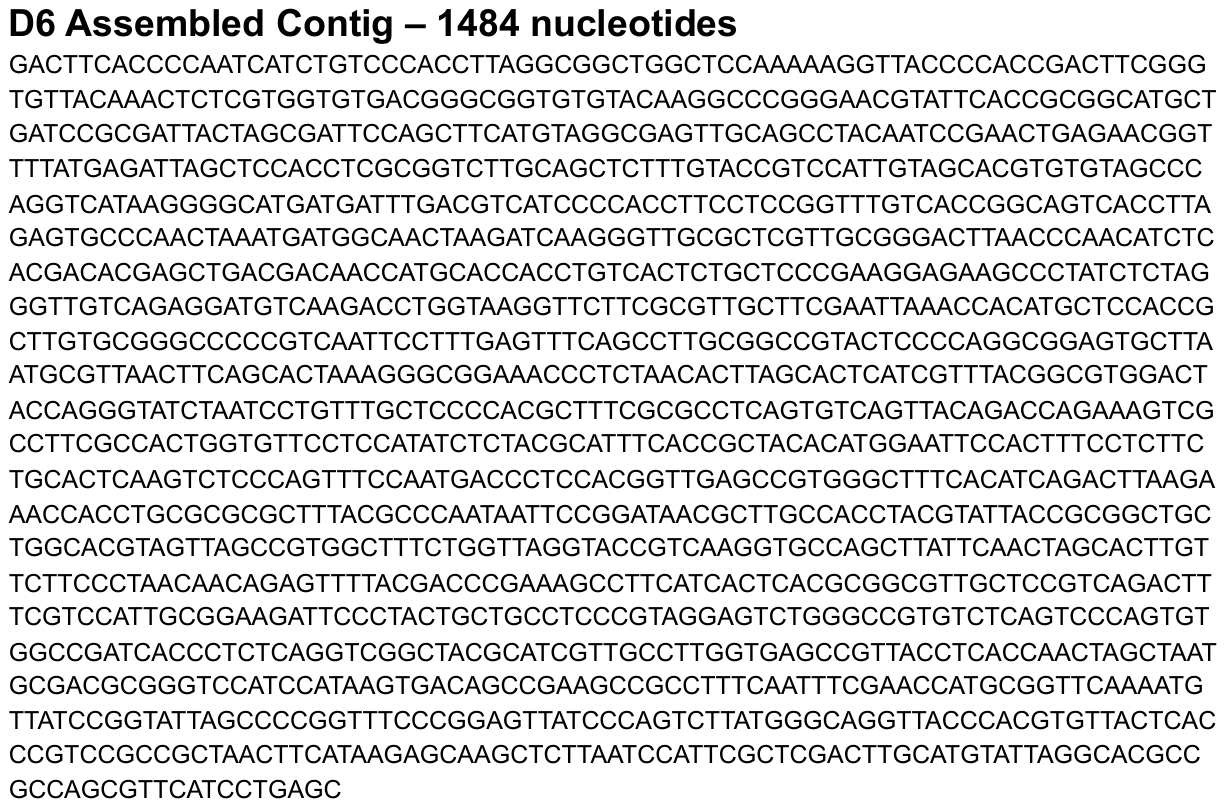

**Figure S24. D6 Assembled Contig – GenBank accession number MT477810**

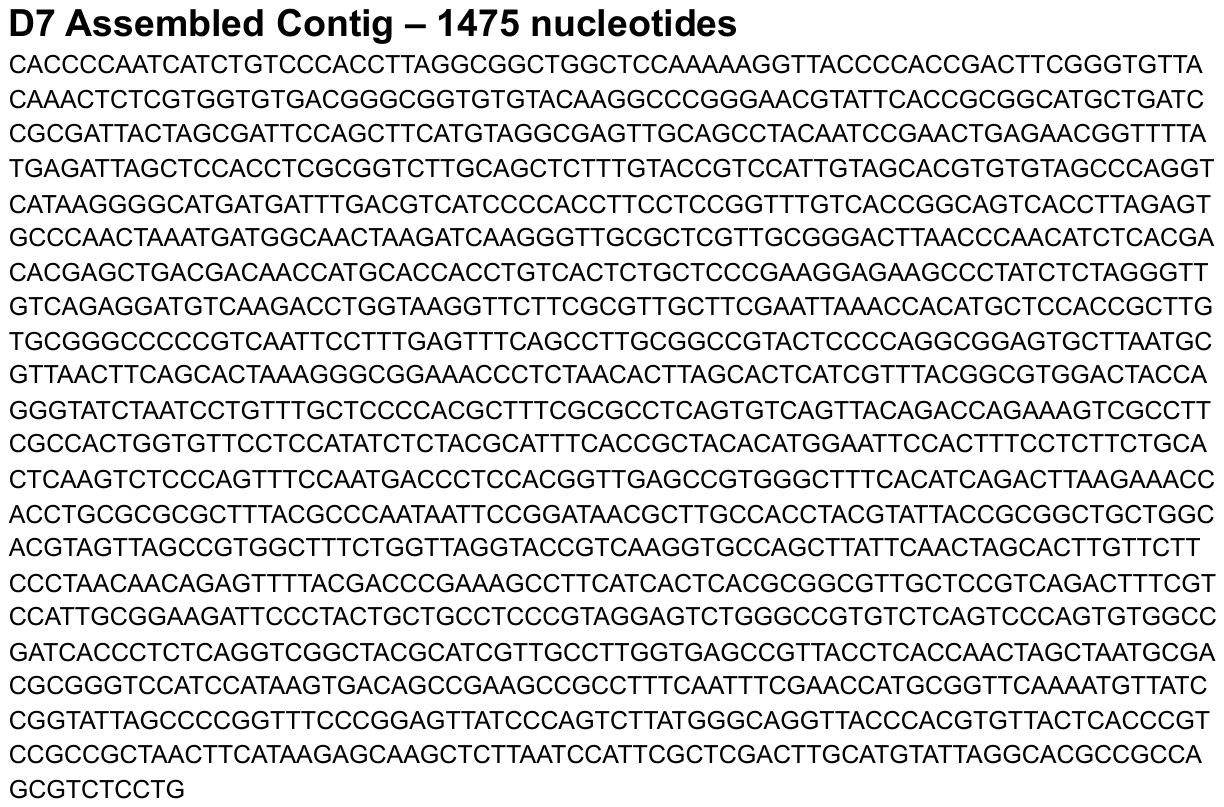

**Figure S25. D7 Isolate assembled contig GenBank accession number MT477812**

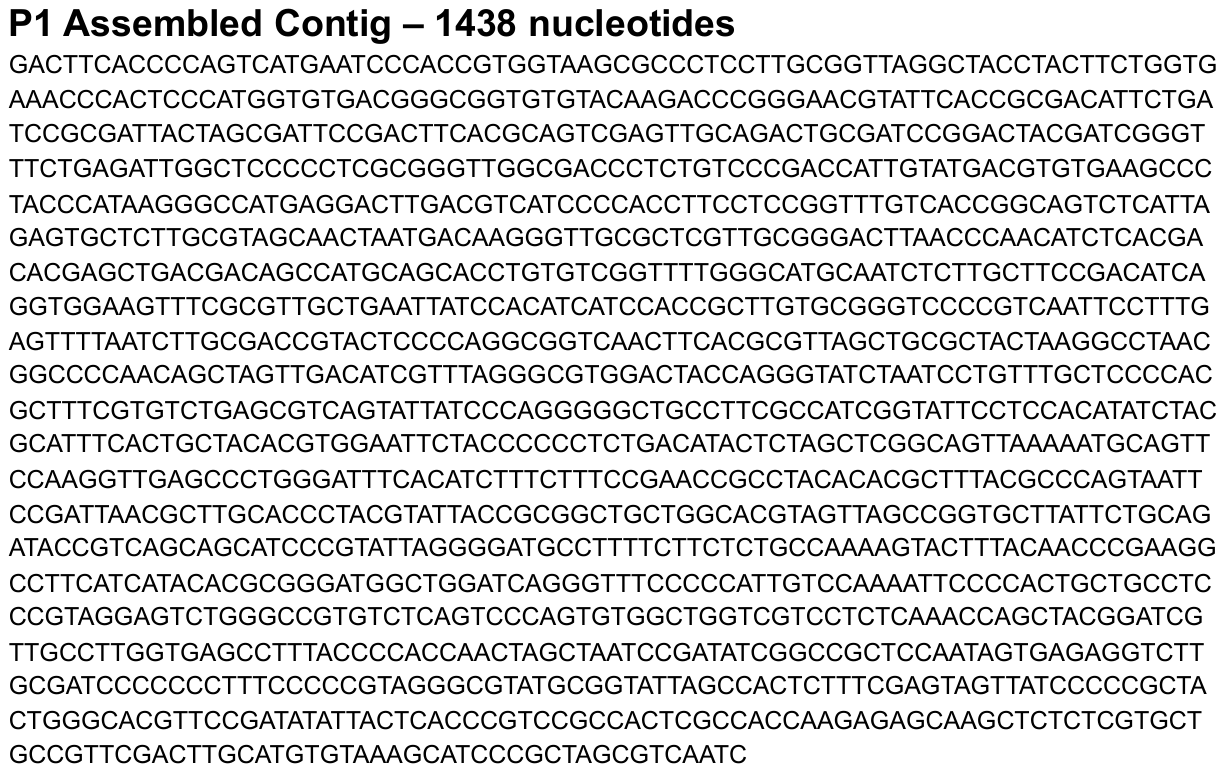

**Figure S26. P1 Isolate assembled contig GenBank accession number MT477813**
